# Viral rewiring of APC/C-CDC20 drives Aurora B hyperubiquitination, mitotic regression and polyploidy

**DOI:** 10.64898/2026.02.13.705602

**Authors:** Shikha Srivastava, Debkanya Sengupta, Yllza Jasiqi, Vanessa R. Povolo, Michael Bauer, Mathieu Hubert, Uta Haselmann, Martin Schneider, Dominic Helm, Urs F. Greber, Vibhu Prasad

## Abstract

Cytokinesis requires elaborate processes in the midbody, including the timely inactivation of Aurora B kinase (AurB) and membrane fission to separate the daughter cells. Here, we show that human adenovirus (AdV) perturbs midbody function, prevents daughter separation and induces cytoplasmic regression of both cancer cells and primary diploid human airway basal cells resulting in polyploidy. Infected cells undergo cleavage furrow regression after midbody formation driven by the viral protein E4orf4 independently of E4orf4 interaction with protein phosphatase 2A. The co-activator of the anaphase-promoting complex/cyclosome (APC/C), CDC20 is upregulated in AdV infection and required for regression. E4orf4 directly interacts with CDC20 and redirects the E3-ubiquitin ligase APC/C-CDC20 towards AurB, promoting hyperubiquitination and premature extraction of AurB from the midbody. Loss of midbody-associated AurB coincides with midbody collapse and furrow regression. Our findings reveal a previously unrecognized APC/C-CDC20-dependent pathway controlling late cytokinesis through AurB extraction from the midbody.

## Introduction

Viruses manipulate host cell cycle machinery to create intracellular environments favorable for replication and persistence^1^. A recurring outcome of such interference is disruption of genome integrity, often manifesting as abnormal ploidy states. Among these, tetraploidy in normal human diploid cells represents a particularly high-risk intermediate, as these cells are intrinsically prone to chromosome mis-segregation, tolerate aneuploidy, and frequently precede malignant transformation^2,3^. Polyploidy broadly denotes cells with DNA content exceeding the diploid state and, depending on cellular context and baseline karyotype, may present as tetraploid or near-tetraploid populations. While oncogenic viruses such as human papillomavirus, Epstein-Barr virus, hepatitis B virus, and Kaposi’s sarcoma-associated herpesvirus exploit polyploidy-promoting mechanisms in human cancers^4–7^, the potential for polyploidy induced during infections not classically considered oncogenic, including adenovirus infection of humans, to generate genomically unstable, pre-malignant cellular reservoirs remains largely unexplored.

Human adenoviruses (AdVs) are large double-stranded DNA viruses that generally cause acute, lytic infections and are efficiently cleared by the immune system. Unlike retroviruses, AdVs do not integrate into the host genome, and productive infection is typically associated with cell death. Along with the lack of clinical evidence for the presence of AdV genetic elements in human cancers^8^, these factors have contributed to their classification as non-oncogenic in humans. Nevertheless, AdVs are capable of long-term persistence in lymphoid cells of the gastro-intestinal tract, tonsils, or lymph nodes, where viral gene expression is restricted and cytolysis is limited^9,10^.

AdV infection has long been associated with profound perturbations of cell cycle control *in vitro* and *in vivo*^11–13^. In cultured cells, AdVs drive quiescent cells into S phase, induce prolonged S-phase arrest, delay mitotic progression, and interfere with cytokinesis, frequently resulting in cells with ≥4N DNA content. These effects are mediated by multiple viral early proteins, including E1A, E1B, and E4 gene products, which override p53- and Rb-dependent checkpoints and uncouple viral DNA replication from host mitotic control^14–18^. Despite evidence of near-tetraploid or polyploid cells arising during AdV infection in cell culture and animal models^12,19,20^, the mechanisms by which AdV induces abnormal ploidy state have remained unclear. As a result, polyploidy has largely been viewed as an abortive byproduct of infection rather than a regulated viral strategy.

Among AdV early proteins, E4orf4 has emerged as a multifunctional regulator of host cell physiology^21^. E4orf4 has been implicated in transcriptional regulation, delayed mitotic progression, induction of cell death, and has been reported to promote the accumulation of near-tetraploid cells in near-diploid lung carcinoma H1299 cells, which subsequently undergo G1 arrest or are cleared via apoptosis^18,22–24^. These activities have been primarily attributed to E4orf4’s interaction with the B55 regulatory subunit of protein phosphatase 2A (PP2A), linking E4orf4 to mitotic control and checkpoint regulation^21,25,26^. However, how E4orf4 induces polyploidy at the cellular level, whether through mitotic slippage, re-replication, or failure of cytokinesis, has not been resolved. Moreover, it remains unclear whether polyploid cells formed during AdV infection are eliminated, arrested, or capable of supporting productive infection.

Although polyploidy is frequently associated with genomic instability and tumor evolution^2,3^, the fate of polyploid cells is highly context-dependent. In immunocompetent tissues, abnormal polyploid cells are often eliminated through cell-intrinsic checkpoints or immune surveillance mechanisms^27^. However, during persistent viral infections characterized by restricted viral gene expression and immune evasion^28,29^, such surveillance may be attenuated. Under these conditions, polyploid cells generated during infection could, in principle, survive long term. Whether AdV-induced polyploid cells are transient, eliminated, or maintained as viable cellular reservoirs has not been investigated.

Here, we identify a defined and conserved mechanism by which AdV infection induces polyploidy through E4orf4-dependent cleavage furrow regression during cytokinesis to generate binucleate cells. Using quantitative live-cell imaging, single-cell fate mapping, proteomics, correlative light electron microscopy, and in vitro binding assays, we show that AdV-infected cells frequently undergo abortive cytokinesis characterized by midbody collapse and regression of daughter cells. This process produces tetraploid cells in normal human diploid cells and higher-order polyploid cells in cancer cells. We demonstrate that this process is conserved in different AdV serotypes, occurs during both acute and persistent infection, and is driven by E4orf4 independently of PP2A binding.

Mechanistically, we identify the anaphase-promoting complex co-activator CDC20 as a direct E4orf4 interactor required for cleavage furrow regression. AdV infection enhances CDC20 levels and prolongs CDC20 residence at the midbody, where E4orf4 redirects the E3-ubiquitin ligase anaphase promoting complex or cyclosome (APC/C) with its adaptor CDC20 activity toward hyperubiquitination and premature extraction of Aurora B kinase (AurB), a key regulator of cytokinesis and abscission^30,31^. Disruption of AurB-dependent midbody integrity results in cleavage furrow regression rather than successful abscission. Importantly, the resulting polyploid cells remain viable, undergo G1/S phase arrest similar to normally dividing cells, and support robust viral gene expression. Together, our findings reveal cytokinesis as a target for AdV interference and uncover a mechanism by which E4orf4 hijacks APC/C-CDC20 signaling to induce polyploidy. These results challenge the view that polyploidy during AdV infection is merely an incidental consequence of infection and provide a framework for understanding how the rewiring of cytokinesis affects polyploidy with enhanced virus output.

## Results

### AdV infection induces cytokinetic furrow regression in acute and persistent infection in vitro independent of PP2A

To systematically characterize how polyploid cells arise during AdV infection, we performed long-term time-lapse imaging of HeLa H2B-mCherry cells infected at high multiplicity of infection (moi=10) to maximize the probability that each tracked cell was infected. As shown in **Fig. 1A**, infected cells progressed through late mitosis and underwent cleavage furrow ingression with clear midbody establishment. However, instead of completing abscission, the cytokinetic bridge collapsed and the two daughter cell bodies regressed back into a single polyploid cell. The two daughter nuclei subsequently formed a compact “congressed” nuclear state within the shared cytoplasm.

**Figure 1.**
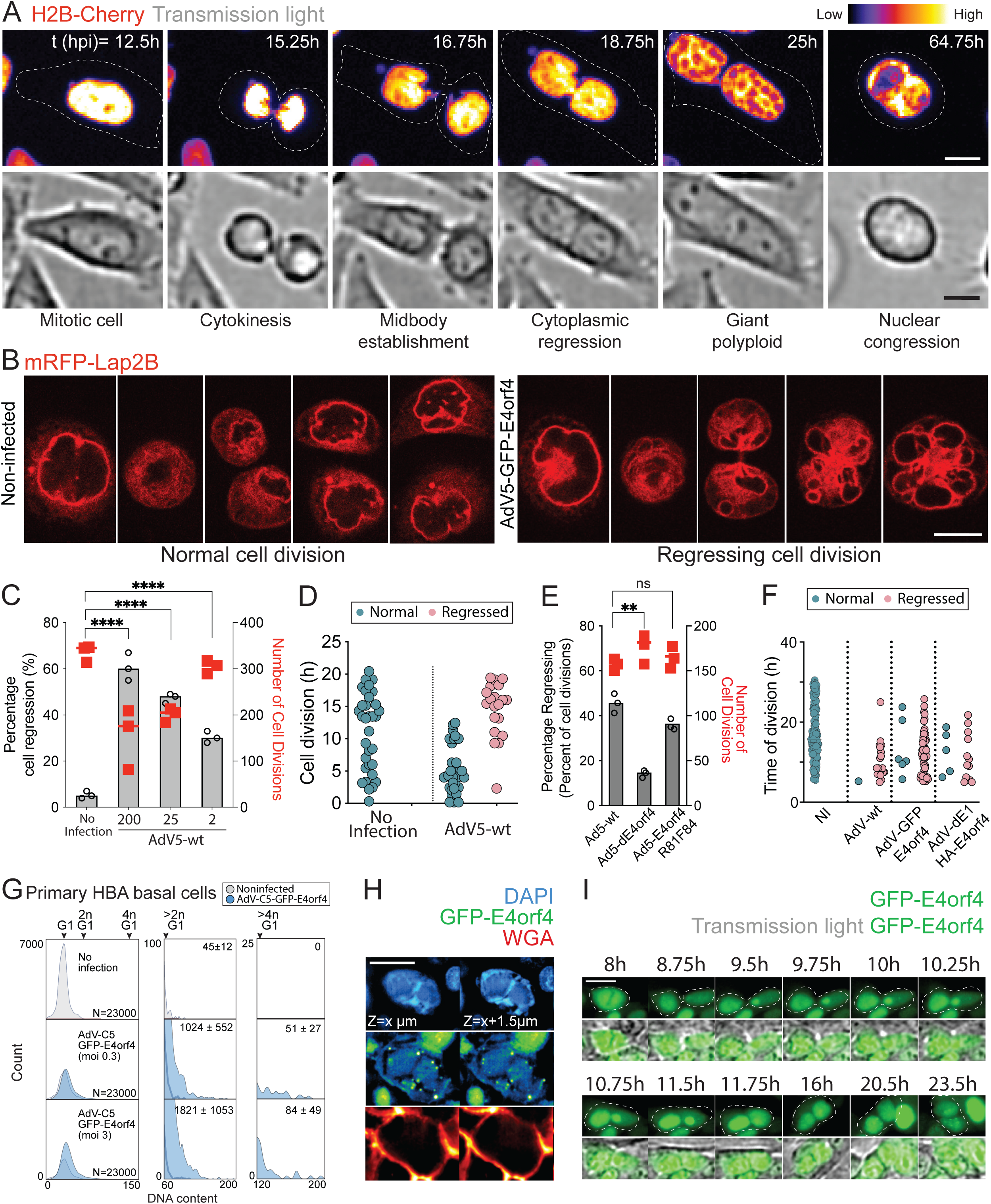
AdV infection induces cleavage furrow regression and polyploidy via the viral factor E4orf4. (A) Cleavage furrow regression in AdV-infected mitotic cells leads to polyploidy. Representative time-lapse images of an AdV-C2-wt-infected HeLa-H2B-mCherry cell undergoing mitosis, showing cytokinesis, cleavage furrow ingression, midbody formation, subsequent furrow regression, and nuclear congression into a single polyploid nucleus. Scale bar, 10 µm. (B) Nuclear envelope remodeling in AdV-induced polyploid cells. Representative confocal images of HeLa cells expressing mRFP-LAP2B show smooth nuclear envelope morphology in non-infected cells, whereas regressed cells generated during AdV infection display pronounced nuclear envelope distortions, including lobulation and membrane invaginations. Scale bar, 10 µm. (C) Viral dose-dependent induction of cleavage furrow regression following AdV5-wt infection. Grey bars indicate the percentage of regressed divisions, and red symbols denote the total number of cell divisions analyzed per condition. Significance was calculated using ordinary one-way ANOVA. **** = p<0.0001. (D) Temporal accumulation of regressed divisions during lytic AdV5-wt infection. Single-cell division times are shown for normal and regressed divisions in non-infected and infected cells. (E) E4orf4 is required for efficient cleavage furrow regression independently of its PP2A-binding function. Deletion of E4orf4 markedly reduces regression frequency, whereas the PP2A-binding-deficient mutant AdV (AdV-C5-E4orf4-R81F84A) retains regression activity. Significance was calculated using one-way ANOVA. ** = p<0.01, ns = p>0.05. (F) E4orf4 expression is sufficient to induce cleavage furrow regression in the absence of productive infection. Time-to-division analysis shows increased regression events upon expression of GFP- or HA-tagged E4orf4 compared with non-infected and vector controls. (G) DNA content analysis of primary human bronchial airway (HBA) basal cells mock-treated or infected with AdV-C5-GFP-E4orf4 at low (MOI 0.3) or moderate (MOI 3) multiplicity. Cells were infected for 48h, followed by fixation, fluorescent microscopy and quantification. Representative histograms show DNA content distributions with G1 peaks, 2NG1, 4NG1, and >4NG1 populations. Quantification indicates a dose-dependent increase in 4N and >4N G1 cells upon infection. Mean ± SD from 3 independent experiments is shown. N = 23,000 nuclei per condition. (H) Confocal imaging of AdV-C5-GFP-E4orf4-infected primary HBA basal cells stained with Hoechst 33342 (DNA) and wheat germ agglutinin (WGA) to delineate cell boundaries. Z-sections illustrate enlarged, irregular nuclei within single-cell boundaries, consistent with polyploidization following cytokinesis failure. Scale bar, 10µm. (I) Time-lapse imaging of AdV-C5-GFP-E4orf4-infected primary HBA basal cells showing progression through mitosis, followed by cleavage furrow regression, binucleated cells. GFP-E4orf4 signal marks infected cells. Dashed outlines indicate cell boundaries and time is shown as hours post infection. Scale bar, 10µm.

To determine whether these congressed nuclei truly fused into one nucleus or remained as two nuclei in close apposition, we imaged HeLa cells stably expressing mRFP-LAP2β (lamina-associated polypeptide 2β), an inner nuclear membrane marker^30^. In regressed cells, the LAP2β signal revealed that nuclear envelopes remained largely distinct but were closely clustered into a single rounded structure with a characteristic lobulated or “flower-like” morphology (**Fig. 1B** and movie S1), consistent with nuclear congression rather than complete nuclear fusion.

We next quantified how frequently AdV infection triggers furrow regression. Early observations indicated substantial heterogeneity in the interval between furrow ingression and regression (fast, intermediate, and slow regressors; **Fig. S1A**), which complicated automated detection of regression events across large datasets. We therefore used a semi-manual scoring strategy, in which infected cells were tracked through division, and each division was annotated as “normal” or “regressed,” with furrow ingression time recorded (**Fig. S1B**). This analysis revealed a strong viral dose dependence, with regression reaching ∼60% of divisions at moi 200, while ∼20-25% of divisions regressed at lower moi (1-2) (**Fig. 1C**). In this system, regression frequency increased notably around ∼10-12 hpi at moi 2 (**Fig. 1D**), consistent with a requirement for viral gene expression. To test this directly, we infected cells with E1-deleted vectors (AdV-C5-dE1-luc or -GFP), which fail to express the viral early gene program because E1 is absent^32^. Under these conditions, we detected no regression (**Fig. S2A**). We also asked whether regression is conserved across AdVs types, and found comparable regression in AdV-B3 infection^33^, supporting the idea that regression reflects a conserved host-virus interaction rather than a serotype-specific feature (**Fig. S2A**).

Near-tetraploidy in AdV infection has previously been linked to the viral E4orf4 protein^24^. Many described E4orf4 activities in cell-cycle control and signaling have been attributed to its interaction with PP2A^21,34^. We therefore tested whether (i) regression requires E4orf4 and (ii) PP2A binding is necessary for regression. An AdV-ΔE4orf4 mutant^35^ was strongly impaired in inducing regression (**Fig. 1E**). Surprisingly, the PP2A-binding-deficient mutant, R81F84A^36^, remained fully competent in triggering regression (**Fig. 1E**). Consistent with this, inhibition of PP2A using okadaic acid^37^ led to only a marginal, non-significant reduction in regression frequency (**Fig. S2B**). Conversely, ectopic E4orf4 expression from a nonreplicating AdV gain-of-function vector (AdV-dE1-E4orf4-HA^24^) was sufficient to induce regression, indicating that E4orf4 is both necessary and sufficient for regression during infection (**Fig 1F**). Similarly, N-terminal GFP-tagged E4orf4 expressing AdV (AdV-C5-GFP-E4orf4^35^) was equally efficient to induce regression. Together, these results support a model in which E4orf4 drives AdV-induced furrow regression largely independently of PP2A binding.

Because AdVs commonly persist in hosts under interferon pressure^38–40^, we asked whether regression is restricted to acute lytic infection or can also occur in an interferon-conditioned persistence model *in vitro*. We performed time-lapse imaging of AdV-infected diploid HDF-TERT cells maintained in a persistently infected state for 18 days under continuous IFNg treatment. Although IFNg reduced proliferation^41^, we observed rare but clear cases of furrow regression in infected cells (an example shown in **Fig. S1C**), whereas no such events were detected in non-infected controls. While this remains qualitative, it provides proof-of-concept that furrow regression can occur under interferon-conditioned persistent infection.

To assess whether AdV-induced cleavage furrow regression and polyploidy also occur in non-transformed cells, we examined primary human bronchial airway (HBA) basal cells derived from healthy donors and infected them with AdV-C5-GFP-E4orf4. As expected for primary cells with low proliferative capacity, DNA content analysis of uninfected cultures revealed a dominant G1 peak with minimal representation of G2/M cells. AdV infection was efficient in these cells, consistent with previous reports^42^, and infection resulted in a clear redistribution of DNA content specifically in GFP-E4orf4-positive cells. Infected cultures exhibited an increased population within the 2N G1-equivalent region, which likely reflects a mixture of low proliferating G2/M cells and tetraploid cells arrested in G1, as well as the appearance of a distinct 4N or higher G1 population that was absent from non-infected controls and were all infected (**Fig. 1G**). High-resolution imaging further identified binucleated cells exclusively within the infected population (**Fig. 1H**). Although live-cell imaging of primary basal cells was technically challenging due to their high migratory behavior (**Fig. S2C**), we were able to capture individual infected cells displaying binucleation (**Fig S2D**) and, in rare cases, a dividing cell undergoing cytokinesis followed by cleavage furrow regression (**Fig. 1I**). Together, these observations demonstrate that AdV-induced cytokinetic failure and near-tetraploidy are not restricted to transformed or polyploid cancer cell lines but can also occur in primary human diploid cells.

### AdV-induced polyploids are viable and support elevated late viral gene expression

Prior work suggested that near-tetraploids generated during AdV infection may die through E4orf4-dependent apoptosis^24^. To directly assess viability and post-mitotic cell-cycle progression of regressed cells, we combined time-lapse imaging with cell-cycle staging using HeLa FUCCI cells^43^. FUCCI reporters distinguish G1 (Cdt1-positive) from S/G2/M (Geminin-positive) states, enabling classification of cell-cycle progression in individual tracked cells, in live or fixed imaging. Time-lapse imaging in AdV-infected HeLa FUCCI cells revealed efficient S and G2/M arrest in infected cells (**Fig. S3A-B**), and that regressed divisions produced viable polyploid cells that re-entered interphase rather than rapidly dying (**Fig. 2A**). Importantly, regression did not produce a distinct FUCCI endpoint compared to non-regressing infected divisions, i.e., both populations accumulated largely at G1/S, consistent with the characteristic AdV-imposed cell-cycle state (**Fig. 2B**)^11,16^. These data argue that regression produces viable infected cells rather than terminally damaged byproducts.

**Figure 2.**
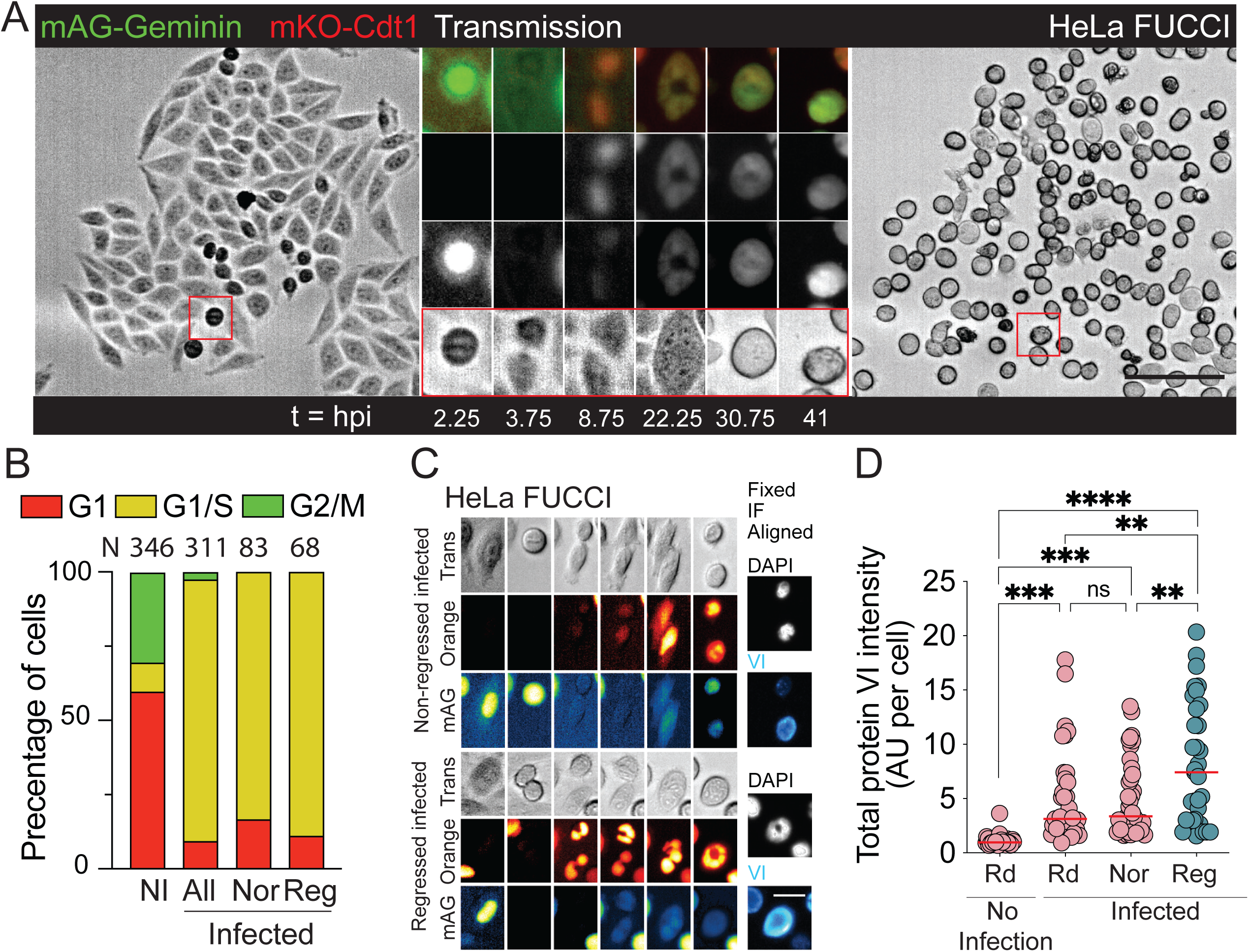
AdV-induced polyploid cells remain viable and support enhanced viral late protein expression. (A) Polyploid cells generated by cleavage furrow regression during AdV infection remain viable and progress through the cell cycle. Time-lapse imaging of AdV-infected HeLa FUCCI cells shows that regressed cells do not undergo cell death but instead complete mitosis, re-enter interphase, and reach the G1/S boundary, similar to non-regressing infected cells. Time shown as hours post infection. Scale bar, 100 µm. (B) Quantification of cell cycle distribution in non-infected (NI) and AdV-infected HeLa FUCCI cells 50 h post infection. Stacked bar plots show the proportion of cells in G1, G1/S, and G2/M phases, with total cell numbers (N) indicated above each condition. Nor: not regressed, Reg: regressed. (C) Correlative time-lapse and fixed-cell immunofluorescence analysis of viral late protein expression in regressed versus normally dividing cells. HeLa FUCCI cells infected with AdV-C5-wt (MOI = 5) were tracked by live imaging to identify regressing and non-regressing divisions, followed by fixation and immunostaining for the viral late protein VI. Fixed-cell images were computationally aligned to the final time point of the time-lapse sequence to assign protein VI intensity to individual tracked cells. Scale bar, 20 µm. (D) Quantification of integrated viral protein VI intensity per cell for non-infected cells, infected non-regressing cells, infected regressing cells, and randomly selected infected cells. Regressed cells show significantly elevated levels of viral late protein expression compared to non-regressing infected cells. Significance was calculated using ordinary one-way ANOVA. ** = p<0.01, *** = p<0.001, **** = p<0.0001. Rd: random; Nor: not regressed, Reg: regressed.

We next asked whether regressed polyploid cells are productively infected. To do this, we performed correlative live-to-fixed imaging. In this setup, regressing and non-regressing divisions were first identified by time-lapse, cells were then fixed and stained for the late viral protein VI, and fixed images were computationally aligned to the last live time point to assign protein VI intensity to each tracked cell (**Fig. 2C**). Regressed cells accumulated significantly higher protein VI signal compared with non-regressing infected cells and randomly sampled infected cells (**Fig. 2D**). This is consistent with the possibility that increased cell size or altered physiology in polyploids supports enhanced late viral protein output. Together, these data suggest that E4orf4-dependent polyploids are viable and are associated with efficient establishment of late viral gene expression.

### CDC20 is a novel direct interactor of E4orf4 required for AdV-induced furrow regression

Since E4orf4-driven regression was largely PP2A-independent (**Fig. 1E; S2B**), we sought alternative host pathways through which E4orf4 might act. Since E4orf4 localizes to both nucleus and cytoplasm^21,44,45^, and furrow regression is a late mitotic or cytoplasmic event, we focused on cytoplasmic E4orf4-associated factors. We therefore performed nucleo-cytoplasmic fractionation of mock- and AdV-C5-GFP-E4orf4-infected cells, followed by GFP-Trap affinity enrichment and mass spectrometry (AP-MS) (**Fig. 3A**). Fractionation quality and GFP-E4orf4 distribution were validated by immunoblotting (**Fig. 3B; S4A**). Comparative analysis identified 59 host and several AdV proteins enriched in the infected cytoplasmic E4orf4 fraction with fold-enrichment of two-folds or higher (**Fig. 3C; S4B; Table S1**). The dataset included several previously reported E4orf4-associated factors, including several subunits of PP2A, GIGYF2, eukaryotic initiation factors isoforms (EIF4E2, EIF3A, EIF5A, EIF3F), CNNM4, and caspase 8^46,47^, supporting technical validity. Among the novel interaction candidates, CDC20 was particularly notable given its established function as APC/C co-activator and its central role in controlling mitotic progression and exit^48,49^. CDC20 has been indirectly linked to E4orf4 through PP2A-dependent pathways^50^, making it plausible that CDC20 could mediate the PP2A-independent mitotic phenotype identified in this study.

**Figure 3.**
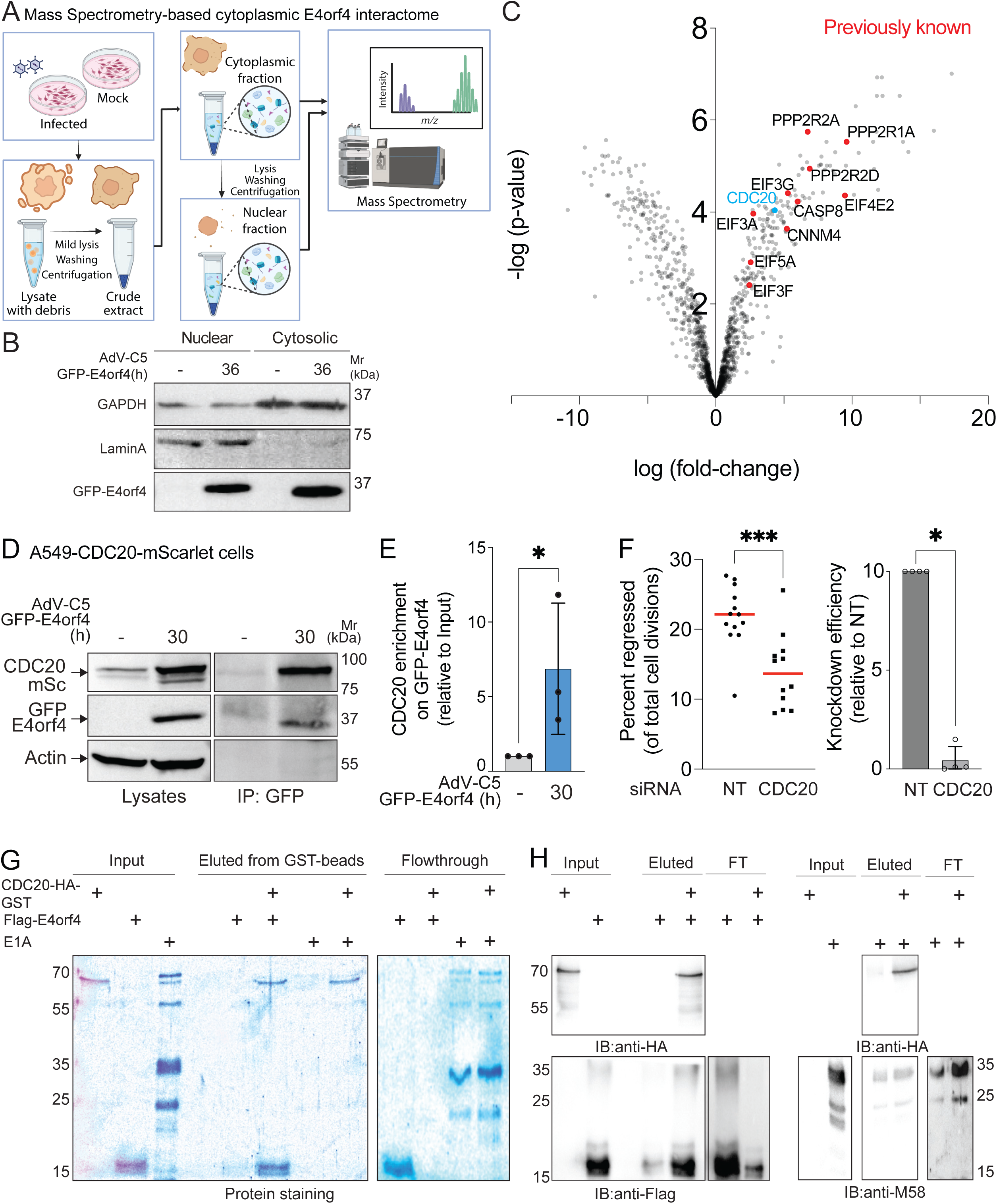
CDC20 is a novel direct E4orf4 interactor required for AdV-induced cleavage furrow regression and polyploidy. (A) Schematic of the workflow used to define the nucleo-cytoplasmic E4orf4 interactome by affinity purification-mass spectrometry. Mock- or AdV-C5-GFP-E4orf4-infected cells were fractionated into nuclear and cytoplasmic compartments, followed by GFP-Trap–based affinity enrichment and mass spectrometric identification of interacting proteins. (B) Immunoblot analysis validating nuclear and cytoplasmic fractionation during AdV-C5-GFP-E4orf4 infection of HeLa cells. GAPDH and Lamin A serve as cytoplasmic and nuclear markers, respectively, confirming enrichment of GFP-E4orf4 in the cytoplasmic fraction. (C) Volcano plot showing proteins enriched in cytoplasmic versus nuclear GFP-E4orf4 interactomes identified by mass spectrometry. Previously reported E4orf4 interactors are highlighted, with CDC20 emerging as a significantly enriched cytoplasmic binding partner. (D) Co-immunoprecipitation of CDC20 with E4orf4 in infected cells. A549 cells expressing CDC20-mScarlet were infected with AdV-C5-GFP-E4orf4, and GFP-Trap immunoprecipitates were analyzed by immunoblotting, demonstrating enrichment of CDC20 in GFP-E4orf4 complexes. (E) Quantification of CDC20 enrichment in GFP-E4orf4 immunoprecipitates relative to input, showing significant association upon AdV infection. (F) Quantification of cleavage furrow regression following CDC20 depletion with RNAi. Time-lapse imaging of AdV-C5-wt–infected A549-Sec61B-GFP cells transfected with non-targeting (NT) or CDC20-specific siRNA shows a significant reduction in the percentage of regressed divisions upon CDC20 knockdown. Each point represents approximately 50 cell divisions, in total 600 cell divisions each condition. Right panel shows reduction in CDC20 mRNA levels following siRNA transfection using quantitative RT-PCR. (G) In vitro GST pull-down assay demonstrating direct binding between CDC20 and E4orf4. GST-CDC20 immobilized on glutathione beads was incubated with purified FLAG–E4orf4 and full-length E1A, followed by SDS-PAGE analysis of input, eluate, and flow-through fractions. (H) Immunoblot validation of GST pull-down assays shown in (G), probing for CDC20 (anti-HA), E4orf4 (anti-FLAG), and E1A (anti-M58), confirming a direct and specific interaction between CDC20 and E4orf4.

We first validated CDC20 association with E4orf4 in infected cells. As endogenous CDC20 is low and cell-cycle regulated^48^, we generated A549 cells stably expressing CDC20-mScarlet and infected them with AdV-C5-GFP-E4orf4. GFP-Trap pulldown showed robust CDC20 enrichment in GFP-E4orf4 complexes (**Fig. 3D**), also relative to input (**Fig. 3E**). We then tested whether CDC20 is functionally required for AdV-induced regression. Time-lapse analysis showed that RNAi depletion of CDC20 significantly reduced regression frequency (**Fig. 3F**), indicating that CDC20 contributes to the regression.

We next asked whether E4orf4 binds CDC20 directly or via a bridging factor such as APC/C. Notably, APC/C subunits were not detected in our cytoplasmic E4orf4 interactome (**Fig. 3C**). We therefore performed *in vitro* binding assays with recombinant proteins. Recombinant CDC20 was expressed and purified as an N-terminal IDR-deleted construct to improve solubility^51^, while His-Flag-E4orf4 was purified in parallel, and an unrelated AdV protein fragment (E1A-WT) was included as a specificity control (**Fig. S5**). GST pull-down assays showed strong enrichment of E4orf4 on CDC20-immobilized beads, with near-complete depletion of E4orf4 from the flowthrough, whereas E1A-WT did not bind CDC20 (**Fig. 3G**). Immunoblotting confirmed specificity using anti-HA (CDC20), anti-Flag (E4orf4), and anti-M58 (E1A) antibodies confirmed interaction specificity (**Fig. 3H**). Together, these data identify CDC20 as a direct E4orf4 interactor and a host factor required for efficient AdV-induced furrow regression.

### CDC20 levels are elevated during AdV infection and cytokinesis of infected cells

We next asked how CDC20-E4orf4 interaction could promote regression. CDC20 functions as a co-activator of APC/C, recruiting substrates for ubiquitylation and proteasome-mediated turnover^48^. CDC20 itself is under tight cell-cycle control, including APC/C-dependent regulation through CDH1^52,53^. Using HeLa FUCCI-CA cells, which allow for sharper transition and separation between G2 and M phases^54^, we investigated CDC20 levels in mock and AdV-infected cells. In uninfected cells, CDC20 peaks in mitosis and declines rapidly during spindle assembly checkpoint at the metaphase-to-anaphase transition^55^, reaching low levels in G1. Consistent with this, endogenous CDC20 was highest in G2/M FUCCI-CA cells and lowest in G1 (**Fig. 4A-B**).

**Figure 4.**
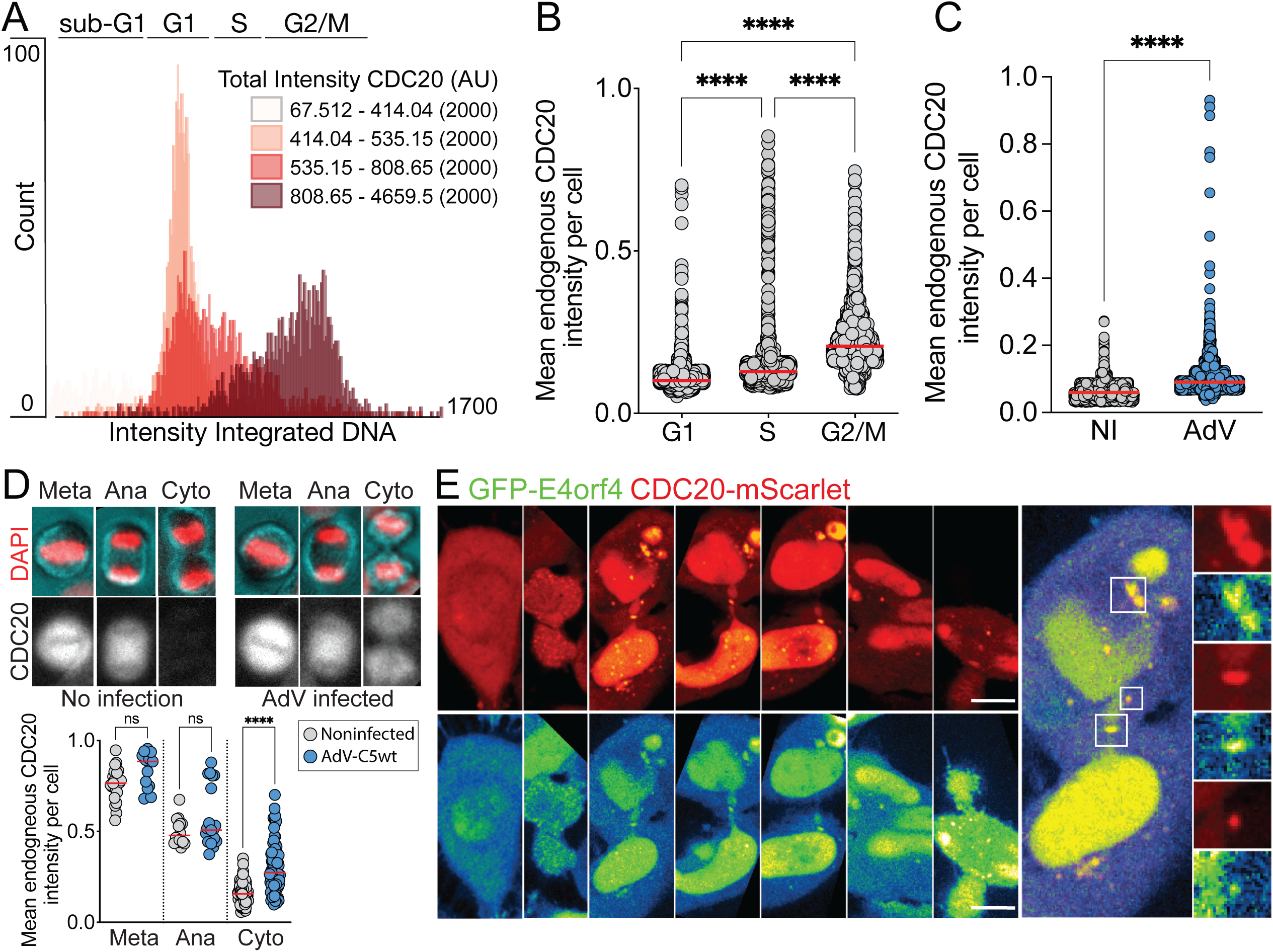
CDC20 levels are elevated in AdV-infected cells and stabilized during cytokinesis. (A) Cell cycle-stage-specific distribution of endogenous CDC20 in HeLa FUCCI cells. Asynchronously cycling HeLa FUCCI cells were fixed and immunostained with an anti-CDC20 antibody. Cells were classified into sub-G1, G1, S, and G2/M phases using a semi-supervised machine-learning classifier (CellProfiler) based on DNA content and FUCCI signals, and CDC20 intensities were mapped onto DNA content-based cell cycle plots. Equal numbers of cells (n = 2,000 per bin) were grouped by increasing CDC20 intensity. (B) Quantification of mean CDC20 intensity per cell across cell cycle stages. Scatter plots show single-cell CDC20 intensities in G1, S, and G2/M phases, revealing highest CDC20 abundance in G2/M cells. Red lines indicate median values; Significance was calculated using ordinary one-way ANOVA. **** = p<0.0001. (C) CDC20 abundance is increased in adenovirus-infected G2/M cells. HeLa FUCCI cells were mock-treated or infected with AdV-C5-wt, synchronized in G2/M by double-thymidine block, fixed, immunostained for CDC20, and quantified as in (A). Scatter plots show increased CDC20 intensity per cell upon AdV infection. Significance was calculated using ordinary one-way ANOVA. **** = p<0.0001. (D) CDC20 stabilization in cytokinetic cells during adenovirus infection. Mitotic metaphase, anaphase, and cytokinetic cells were identified in mock- and AdV-infected A549 cells using the classification strategy described in (A). Quantification of CDC20 intensity shows selective stabilization of CDC20 in cytokinetic cells following AdV infection. ns, not significant; Significance was calculated using ordinary one-way ANOVA. **** = p<0.0001. (E) Live-cell imaging of CDC20-E4orf4 co-dynamics during cleavage furrow regression. A549 cells expressing CDC20-mScarlet were infected with AdV-C5-GFP-E4orf4 and imaged by spinning-disk confocal microscopy. Representative time-lapse images show co-accumulation of CDC20-mScarlet and GFP-E4orf4 in a cytokinetic cell, coinciding with midbody collapse and cleavage furrow regression. Scale bar, 10µm.

During AdV infection, CDC20 levels were markedly increased. We initially observed this in CDC20-overexpressing A549 cells (**Fig. 3E**), but because AdV infection can induce broad transcriptional programs such as UPR signaling^39^, we verified the effect at the endogenous level. Both, analyses of synchronized and non-synchronized cells showed significantly increased endogenous CDC20 intensity in infected cells (**Fig. 4C; Fig. S6A-B**), in line with ectopic CDC20 expression.

Because CDC20 is degraded after spindle assembly checkpoint activation in late mitosis^55^, we asked whether CDC20 is maintained during cytokinesis of infected cells. Infected cells showed higher CDC20 levels in cytokinesis compared with mock-treated controls, where CDC20 dropped sharply post anaphase (**Fig. 4D**). Live imaging further supported CDC20 presence at cytokinetic bridges in infected cells, where CDC20-mScarlet signal overlapped with GFP-E4orf4 in a regressing division (**Fig. 4E**). These data support the idea that AdV infection not only elevates general CDC20 abundance but importantly also in cytokinesis.

### E4orf4 promotes APC/C-CDC20-dependent Aurora B hyperubiquitination

E4orf4 binding to CDC20 suggested that E4orf4 may hijack APC/C-CDC20 to drive regression by altering ubiquitylation of late mitotic substrates. Alternatively, E4orf4 itself could be an APC/C substrate. To probe whether E4orf4 is proteasome-regulated, we treated infected cells with the proteasome inhibitor MLN9708/ixazomib^56^ and observed strong stabilization of E4orf4 (**Fig. S7A-B**). We then quantified GFP-E4orf4 dynamics during infection by imaging regressing AdV-C5-GFP-E4orf4-infected HeLa H2B-mCherry cells. GFP-E4orf4 displayed pronounced oscillations after its initial appearance, and regression occurred close to peak E4orf4 intensity (**Fig. 5A-B; Fig. S7C**). These data suggest that E4orf4 turnover is temporally coupled to the regression event.

**Figure 5.**
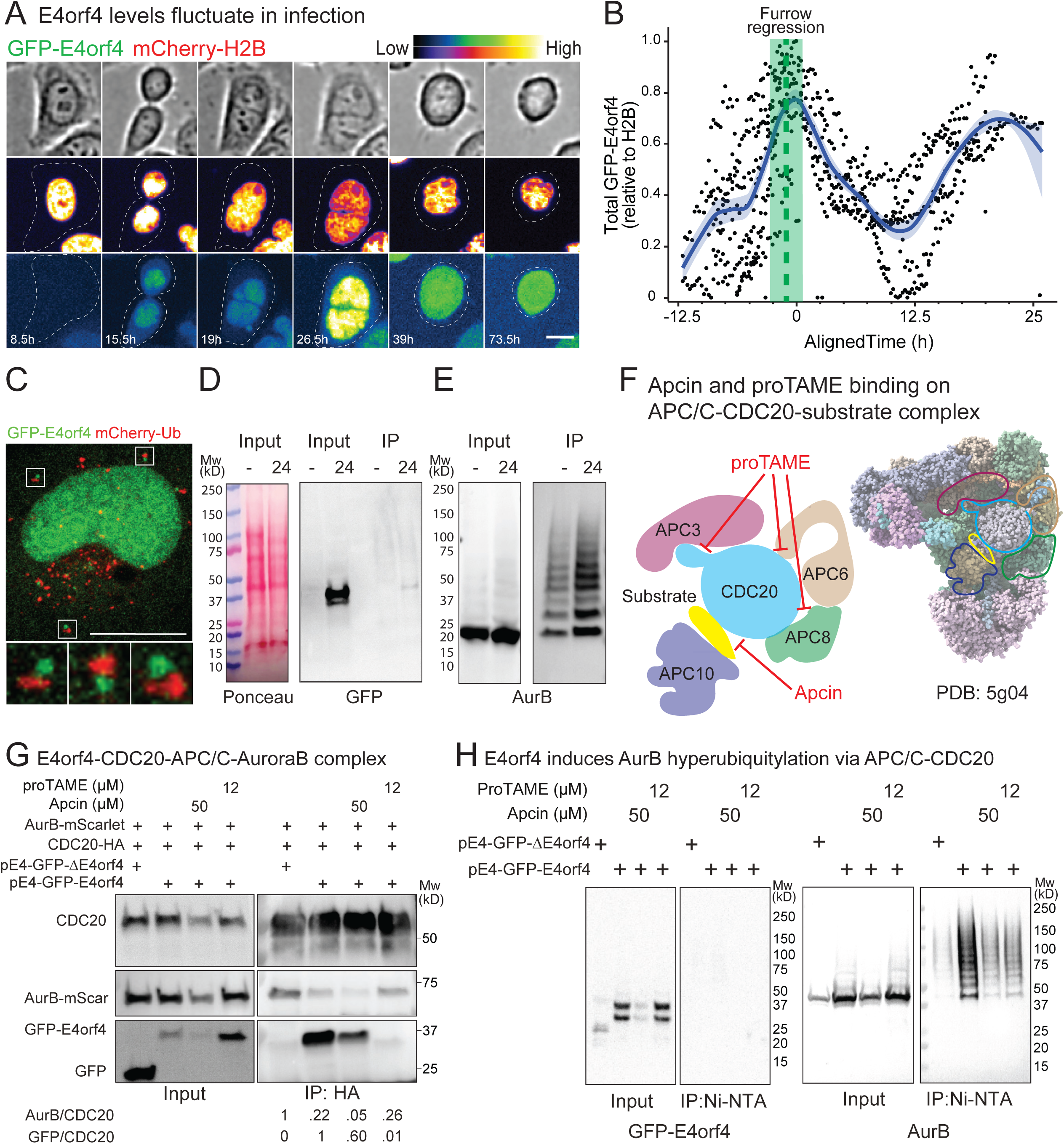
E4orf4 promotes APC/C-CDC20-dependent hyperubiquitination of Aurora B. (A) Live-cell imaging of GFP-E4orf4 dynamics during AdV infection. Representative time-lapse images of HeLa-H2B-mCherry cells infected with AdV-C5-GFP-E4orf4 show fluctuating levels of GFP-E4orf4 over time (shown as hours post infection). Pseudocolor indicates relative fluorescence intensity (low to high). Scale bar, 10 µm. (B) Time-aligned single-cell analysis of GFP–E4orf4 expression dynamics. GFP-E4orf4 intensity traces from individual cells were aligned to the time point of maximal E4orf4 expression, revealing a common oscillatory pattern and a peak coinciding with cleavage furrow regression (green shaded region). (C) Co-localization of GFP-E4orf4 and ubiquitin in infected cells. HeLa cells stably expressing mCherry-ubiquitin were infected with AdV-C5-GFP-E4orf4 for 24 h, fixed, and imaged by confocal microscopy, revealing accumulation of ubiquitin-rich structures in E4orf4-positive cells. Scale bar, 10 µm. (D) E4orf4 itself is not detectably ubiquitylated during adenovirus infection. HEK293T cells transfected with His-tagged ubiquitin were infected with AdV-C5-GFP-E4orf4 for 24 h, lysed under denaturing conditions, and ubiquitylated proteins were enriched by Ni-NTA pulldown. Immunoblotting with anti-GFP antibody shows no high-molecular-weight ubiquitylated E4orf4 species. (E) Aurora B undergoes hyperubiquitination during adenovirus infection. HEK293T cells transfected with His–ubiquitin and AurB-HA were infected and processed as in (D). Ni-NTA pulldown followed by immunoblotting with AurB-specific antibodies reveals extensive high-molecular-weight ubiquitylated Aurora B species. (F) Schematic representation of APC/C-CDC20 inhibition by Apcin and proTAME. Based on the APC/C-CDC20 structure (PDB: 5g04), Apcin blocks the D-box substrate–binding interface of CDC20, whereas proTAME interferes with CDC20 binding to APC/C TPR subunits (APC3, APC6, APC8). Inhibitor positions are schematic and indicate functional inhibition interfaces rather than exact atomic contacts. (G) Formation of an E4orf4-CDC20-APC/C-AurB complex requires an APC/C-bound CDC20 state. HEK293T cells were transfected with plasmids expressing wild-type or ΔE4orf4 adenoviral constructs together with CDC20-HA and AurB-mScarlet and treated with Apcin or proTAME as indicated. CDC20 immunoprecipitates (anti-HA) were analyzed by immunoblotting, demonstrating E4orf4- and APC/C-dependent association of AurB with CDC20. (H) E4orf4 directs APC/C-CDC20 activity toward AurB hyperubiquitination. HEK293T cells were transfected with His–ubiquitin, AurB-HA, and wild-type or ΔE4orf4 constructs, treated with Apcin or proTAME, and briefly exposed to the proteasome inhibitor MLN9708. Ubiquitylated proteins were enriched by Ni-NTA pulldown and immunoblotted, showing E4orf4- and APC/C-CDC20-dependent hyperubiquitination of AurB.

To test whether E4orf4 is directly ubiquitylated, we first examined Ub-mCherry localization in infected cells. GFP-E4orf4 was frequently found near ubiquitin-rich structures but did not strongly colocalize (**Fig. 5C**), consistent with either transient or indirect association of E4orf4 with ubiquitin or ubiquitination of another complex. We then performed denaturing Ni-NTA pulldowns from His-Ub-expressing HEK293T cells infected with AdV-C5-GFP-E4orf4. GFP-E4orf4 was not recovered in the ubiquitylated fraction (**Fig. 5D**), arguing against robust direct ubiquitylation of E4orf4 under these conditions.

We therefore considered that E4orf4 might retarget APC/C-CDC20 to a midbody-associated substrate, whose abnormal ubiquitylation could impair cytokinesis. Aurora B kinase (AurB) is a central master-regulator of cytokinesis coordinating midbody signaling and abscission timing^30,57^. AurB can be regulated by ubiquitylation through APC as a substrate via the co-activator Cdh1, and is capable of binding to CDC20^58^. Using denaturing His-Ub pulldown assays, we detected prominent high-molecular-weight AurB signals in SDS-PAGE from infected cell lysates, consistent with AurB hyperubiquitination (**Fig. 5E**). To assess whether E4orf4 promoted AurB hyperubiquitination independently of other viral factors, we expressed E4orf4 from the viral E4 promoter to approximate an infection-like regulation and avoid toxicity seen from constitutive expression^26^ (**Fig. S8A**). HEK293T cells with stable E1A-expression^59^ were transfected with pE4-GFP-E4orf4 or a matched ΔE4orf4 control plasmids showed robust yet controlled expression of GFP-E4orf4 relative to GFP without toxicity (**Fig. S8B**). We then tested APC/C-CDC20 dependence using Apcin and proTAME^60^, the former inhibiting CDC20 substrate engagement and the latter preventing co-activator binding to APC/C by mimicking the C-term IR-tail of CDC20, hence preventing CDC20 recruitment to APC/C via the TPR lobe involving the subunits APC6/8/3, respectively (**Fig 5F**). CDC20-HA immunoprecipitation revealed robust CDC20-AurB association in controls and formation of E4orf4-CDC20-AurB complex upon E4orf4 expression (**Fig. 5G**). Apcin reduced CDC20-AurB binding but did not substantially disrupt CDC20-E4orf4 association, consistent with AurB behaving as a CDC20-engaged substrate (**Fig. S8C**). In contrast, proTAME strongly reduced CDC20-E4orf4 association, suggesting that E4orf4 preferentially engages CDC20 in an APC/C-bound state or requires an intact IR-tail interaction network. We used the same setup as described in **Fig 5G** in the presence of His-Ub transfection and immunoprecipitation of Ub-proteins via Ni-NTA under denaturing conditions to identify cellular ubiquitylated proteins. Importantly, E4orf4 expression strongly promoted AurB hyperubiquitination, and this was inhibited by both Apcin and proTAME (**Fig. 5H**). Together, these results support a model in which E4orf4 binds CDC20 and associates in a ternary E4orf4-CDC20-AurB complex to redirect APC/C-CDC20 activity toward AurB, leading to AurB hyperubiquitination.

### Hyperubiquitylated Aurora B is prematurely extracted from the midbody, coinciding with midbody collapse and furrow regression

AurB activity restrains abscission, and premature Aurora B inactivation can drive cytokinesis failure and regression^30^. AurB activation is commonly monitored via T232 phosphorylation (pT232). In furrow-ingressed cytokinetic cells infected with AdV-C5-GFP-E4orf4, we detected significantly reduced endogenous AurB and AurB pT232 levels at the midbody region relative to non-infected cells (**Fig. 6A-B**), consistent with premature loss of AurB activity at the cytokinetic bridge.

**Figure 6.**
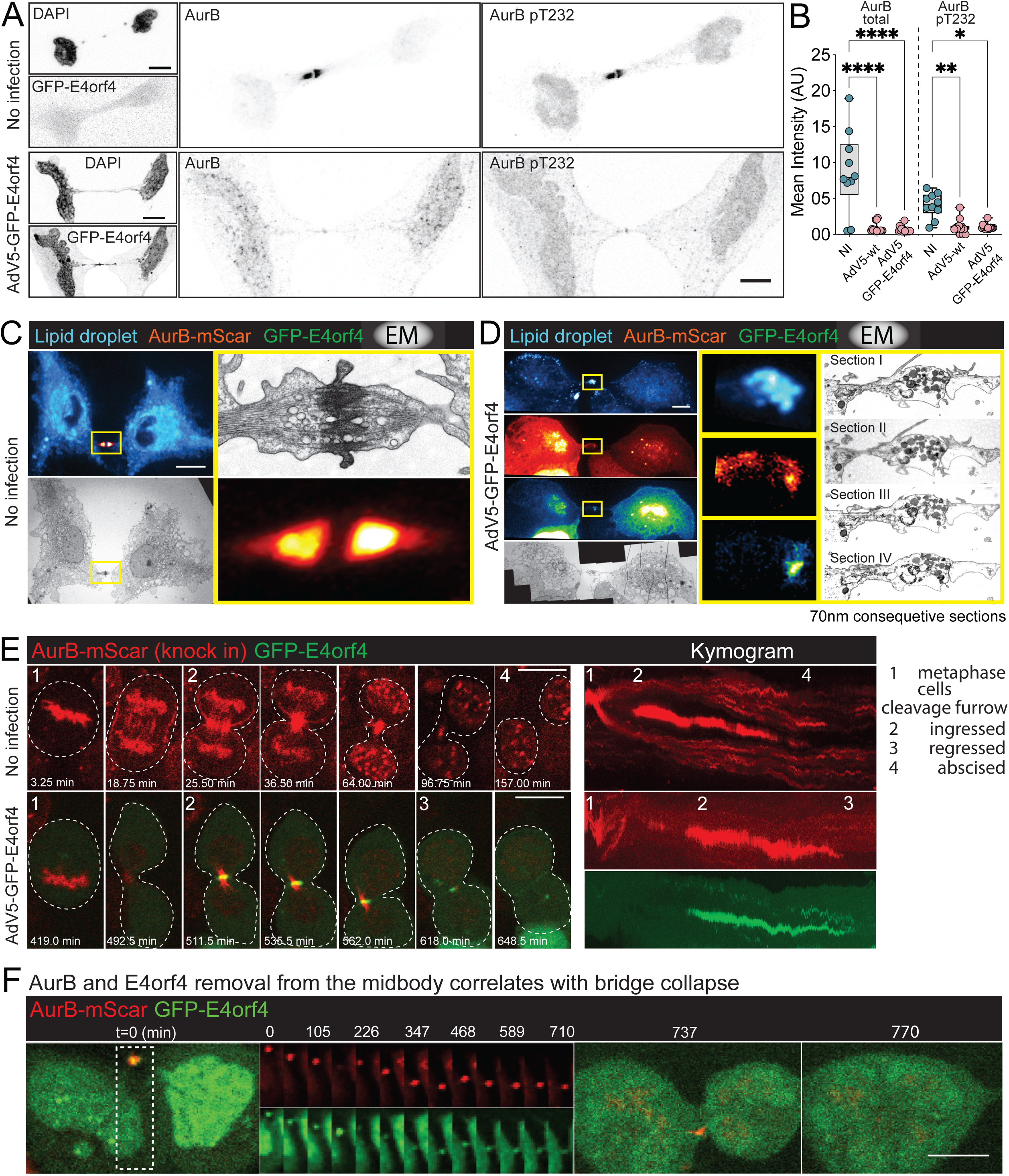
Aurora B is extracted from the midbody prior to cleavage furrow regression in AdV-infected cells. (A) AdV infection reduces endogenous AurB and active AurB (pT232) levels at the midbody of cytokinetic cells. HeLa cells were infected with AdV-C5-GFP-E4orf4 for 24 h, fixed, and immunostained for AurB and AurB pT232. Cytokinetic cells with an ingressed cleavage furrow were identified by confocal microscopy and imaged. Scale bar, 10 µm. (B) Quantification of AurB and AurB pT232 signal intensity at the midbody in mock- and AdV-infected cytokinetic cells shown in (A). Significance was calculated using ordinary one-way ANOVA. **** = p<0.0001. (C–D) AdV infection alters midbody ultrastructure. AurB-mScarlet-expressing A549 cells were seeded on 3.5-mm gridded dishes and mock-treated (C) or infected with AdV-C5-GFP-E4orf4 for 24 h (D). Cells with ingressed cleavage furrows were identified by live imaging, fixed, and processed for correlative light and electron microscopy (CLEM). Fluorescence and electron micrographs were aligned using lipid droplets as fiducial markers (cyan). Yellow boxes indicate regions shown at higher magnification. Serial 70-nm sections are shown for infected cells. Scale bar, 10µm. (E) Endogenously tagged AurB-mScarlet is removed from the midbody prior to cleavage furrow regression in AdV-infected cells. HEK293T cells carrying a C-terminal mScarlet knock-in at the AURKB locus (generated by microhomology-mediated end joining) were infected with AdV-C5-GFP-E4orf4 and imaged by spinning-disk confocal microscopy. Representative time-lapse images show AurB and GFP-E4orf4 localization during cytokinesis in non-infected and infected cells. Kymograms (right) illustrate cleavage furrow ingression in both conditions, followed by abscission in non-infected cells and regression in infected cells. Scale bar, 10µm. (F) Live-cell imaging of AurB-mScarlet–expressing A549 cells showing loss of Aurora B from the midbody preceding cleavage furrow regression in AdV-infected cells. Scale bar, 10 µm.

We next asked whether infected midbodies exhibit ultrastructural abnormalities. Using correlative light-electron microscopy (CLEM), A549 AurB-mScarlet cells were seeded on gridded dishes and imaged live to identify cytokinetic bridges in mock or infected samples. Lipid droplets were used as fiducial markers for correlation (**Fig. 6C-D**). Uninfected midbodies showed typical ultrastructure with electron dense structures, vesicles and coaxial microtubules^61,62^, whereas infected midbodies displayed markedly altered architecture, disturbed bridge organization across serial sections and reduced AurB signal (**Fig. 6D**).

To directly track AurB in living infected cells, we generated HEK293T cells with an endogenous AurB-mScarlet knock-in via MMEJ (ref^63^ **Fig. S9A-B**). In non-infected divisions, AurB exhibited canonical CPC dynamics, forming a stable midbody track through abscission and resolving into a midbody remnant(**Fig. 6E** and movie S2)^30,64–66^. In contrast, infected cells recruited AurB toward the bridge but failed to maintain a confined, stable midbody signal. Instead, AurB signal was lost from the bridge prior to abscission and this loss coincided with furrow regression (**Fig. 6E** and movie S3). In A549 AurB-mScarlet cells, where the midbody morphology is often more extended, AurB removal prior to bridge collapse was particularly clear (**Fig. 6F**). Together, these live imaging data argue that AdV infection promotes premature AurB loss or extraction from the midbody prior to regression.

Given our biochemical evidence that E4orf4 promotes APC/C-CDC20-dependent Aurora B hyperubiquitination (**Fig. 5H**), we next examined ubiquitin dynamics at the midbody. In Ub-GFP/AurB-mScarlet A549 cells, Ub-GFP association with the midbody was transient during normal divisions but became strikingly prolonged in infected cells, with ubiquitin persisting at AurB-positive midbodies until shortly before regression (**Fig. 7A-C** and movie S4-5). This was also observed prior to regression using GFP-E4orf4 presence at the midbody in primary human bronchial airway basal cells (**Fig. S10A**), with Ub-mCherry (**Fig. S10B** and movie S6) or AurB-mScarlet and Ub-BFP-expressing cells (**Fig. S10C** and movie S7).

**Figure 7.**
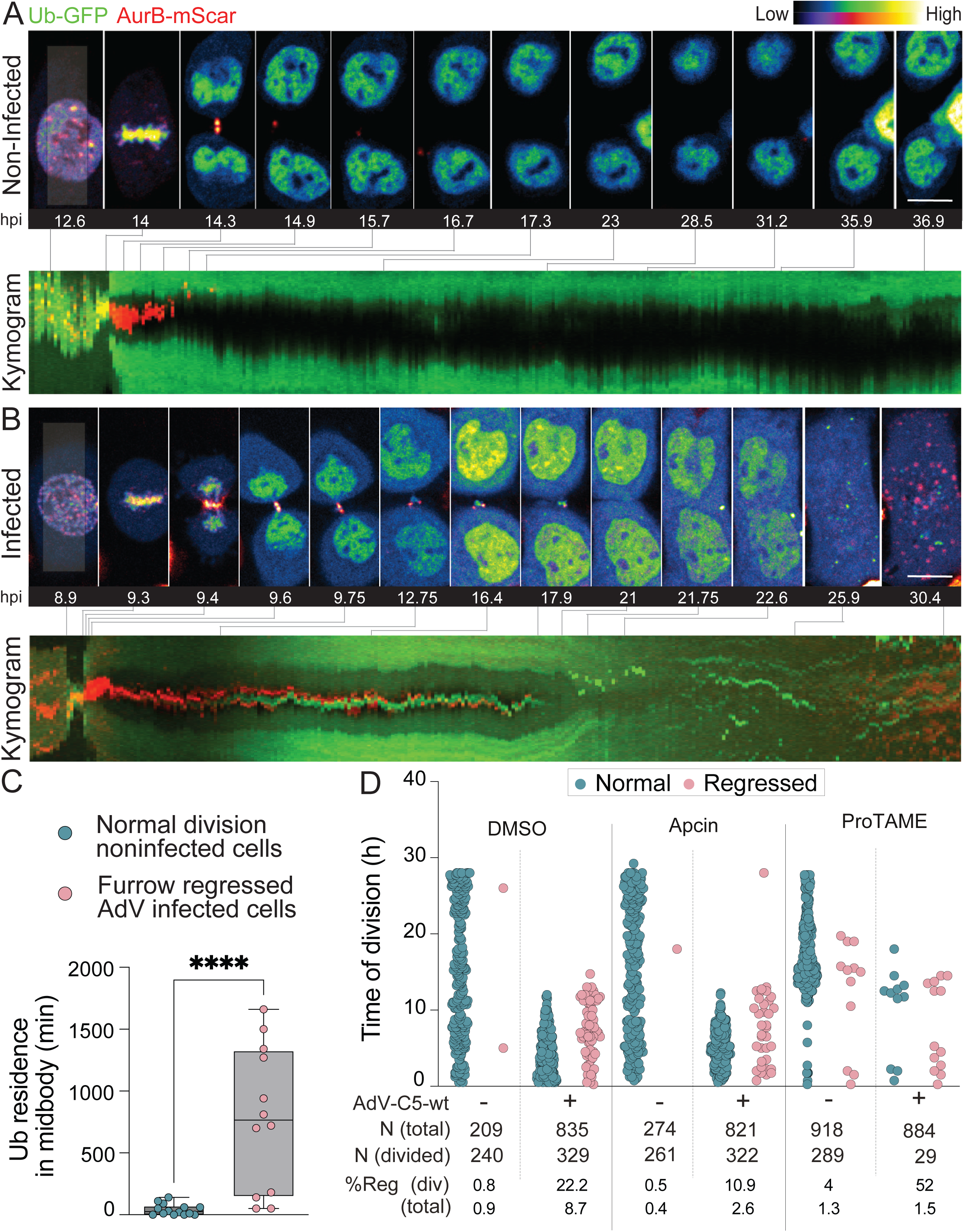
APC/C-CDC20-dependent Aurora B removal drives midbody collapse and polysploidy during AdV infection. (A-B) Prolonged ubiquitin residence at midbody-localized Aurora B during AdV infection. A549 cells expressing Ub-GFP and AurB-mScarlet were mock-treated (A) or infected with AdV-C5-wt (B) and imaged by spinning-disk confocal microscopy. Representative time-lapse sequences show progression from cleavage furrow ingression to abscission in non-infected cells or to furrow regression in infected cells. Corresponding kymograms (bottom) indicate the temporal alignment of individual frames and reveal sustained ubiquitin association with AurB at the midbody in infected cells prior to its removal and subsequent regression. Scale bar, 10µm. (C) Quantification of ubiquitin residence time at the midbody. Box-and-whisker plots show the duration of Ub-GFP persistence at AurB-positive midbodies in normal divisions of non-infected cells and regressed divisions of AdV-infected cells. Each point represents one cell. Significance was calculated using ordinary one-way ANOVA. **** = p<0.0001. (D) APC/C inhibition suppresses adenovirus-induced cleavage furrow regression via distinct mechanisms. A549-Sec61β-GFP cells were mock-treated or infected with AdV-C5-wt and treated with DMSO, Apcin, or proTAME as indicated. Single-cell time-lapse analysis shows the timing and outcome of cell divisions, with normal and regressed divisions plotted separately. Apcin significantly reduces the frequency of cleavage furrow regression, whereas proTAME prevents infected cells from progressing beyond mitosis. Percentages of regressed divisions and numbers of analyzed and divided cells are indicated.

Finally, we tested the impact of APC/C-CDC20 inhibition on suppressing regression in infected cells. To this end, Apcin reduced regression frequency by 50%, whereas proTAME caused infected cells to stall in mitosis and largely prevented progression to regression (**Fig. 7D, S11**). Notably, proTAME also disrupted CDC20-E4orf4 association (**Fig. 5G**), supporting the idea that E4orf4 induction of cleavage furrow regression requires an APC/C-bound CDC20 state. Together, these data support a model in which E4orf4 hijacks APC/C-CDC20 at the midbody to drive excessive AurB ubiquitylation and premature AurB removal, destabilizing the cytokinetic bridge and triggering cleavage furrow regression.

## Discussion

In this study, we identified cleavage furrow regression as a prominent and conserved outcome of AdV-infection in proliferating cells. Using live-cell imaging, we showed that AdV-infected cells progress through furrow ingression and midbody formation but fail to completely abscise the midbody, resulting in midbody collapse and the formation of viable polyploid cells. This phenotype was observed in different AdV serotypes and occurred both in acute lytic infection and in a persistent infection model, indicating that cytokinetic failure is not restricted to high viral loads or cytopathic conditions. While AdV-induced polyploidy has been reported previously^24^, our work provides direct spatiotemporal evidence that polyploids arise majorly through regression of the cleavage furrow rather than failed mitotic entry or mitotic slippage.

Cytokinetic regression is generally considered a rare or pathological outcome, occurring under conditions of chromatin bridges, unresolved DNA damage, or perturbation of abscission checkpoints^30,31^. The significant frequency of regression observed in AdV infected cells, reaching up to 20-25% of divisions at an moi of 1-2, suggests that AdV actively reprograms late mitotic events rather than passively triggers a stress response with pleiotropic effects. Importantly, in our system, polyploid cells generated by AdV-induced cleavage furrow regression are viable, re-enter the cell cycle, and showed efficient viral gene expression, indicating that they escape from terminal elimination by apoptotic mechanisms as previously thought^24^. Polyploidy is widely recognized as an unstable intermediate state that promotes chromosomal instability and facilitates the accumulation of oncogenic mutations^2,3^. In our experimental setup *in vitro*, lytic AdV infection in primary human bronchial airway basal cells and persistent AdV infection in diploid HDF-TERT cells^67^ also showed tetraploid cell formation via cleavage furrow regression, which might be of particular significance given that AdVs display oncogenic potential in experimental mammalian models^68^ and that their ability to persist in diverse human cell types has raised longstanding questions regarding HAdV oncogenicity^69^. In these contexts, viral early gene products can continue to modulate host cell cycle pathways while largely escaping immune detection. We propose that such persistently infected near-tetraploid cells may constitute a previously unrecognized, pre-malignant cellular reservoir generated through virus-driven perturbation of cytokinesis and ploidy control and warrant a deeper mechanistic understanding.

Our data identify the viral protein E4orf4 as a key driver of AdV-induced cleavage furrow regression. Loss-of-function and gain-of-function approaches demonstrate that E4orf4 is both necessary and sufficient to induce regression in dividing cells. Although many previously described E4orf4 functions, including apoptosis induction and cell cycle perturbation, have been attributed to its interaction with PP2A^21,70^, we find that the cytokinetic phenotype described here is independent of PP2A. A PP2A-binding-defective E4orf4 mutant retained regression-inducing activity, and pharmacological PP2A inhibition had only marginal effects. These findings expand the functional repertoire of E4orf4 and indicate that it engages distinct host pathways in different subcellular sites and with different cellular interactors. In particular, our results suggest that E4orf4 exerts a PP2A-independent role during late mitosis and cytokinesis, consistent with earlier observations that E4orf4 localizes to cytoplasmic and membrane-associated structures in addition to the nucleus^21,44,45^. Contrary to earlier suggestions that AdV-induced near-tetraploids are destined for apoptosis^24^, our single-cell analyses demonstrate that regressed cells remain viable, reach G1/S phase arrested state, and robustly express late viral proteins. Indeed, polyploid cells exhibited higher levels of later viral protein VI compared to non-regressing or randomly selected infected cells, suggesting that increased cell size and cellular or viral genomic content may favor viral gene expression. This observation aligns with the general principle that viruses benefit from enlarged biosynthetic capacity and relaxed cell cycle control^1,71^.

By focusing on cytoplasmic E4orf4 complexes, we identify CDC20 as a novel and functionally important E4orf4 interactor. CDC20 is best known as a co-activator of the anaphase-promoting complex/cyclosome (APC/C), where it controls the timely degradation of mitotic regulators to ensure orderly mitotic exit^72^. Although an indirect connection between E4orf4 and CDC20 via PP2A had been proposed previously^50^, our biochemical reconstitution experiments demonstrate a direct interaction between E4orf4 and CDC20. Our data further clarify why E4orf4-driven cleavage furrow regression occurs independently of PP2A, despite PP2A’s established role in activating CDC20^73^. PP2A dephosphorylates CDC20 to promote its canonical substrate engagement with APC/C^73^, yet AdV-induced regression under PP2A inhibition (via okadaic acid) or attenuated APC/C recruitment (via E4orf4 PP2A-interaction mutant) persists. This is consistent with PP2A’s preference to dephosphorylate phosphothreonine residues to promote binding of threonine-rich degrons to CDC20 substrate-recognition domains^74–76^. In contrast, E4orf4 associates with CDC20 outside these substrate-binding interfaces, as shown by the insensitivity of the complex to Apcin. Consequently, PP2A-mediated dephosphorylation is unlikely to influence E4orf4-CDC20 complex formation. Functionally, CDC20 is required for AdV-induced regression, as its depletion significantly reduces regression frequency. We show that AdV infection enhances CDC20 expression and allows its persistence into cytokinesis, a stage at which CDC20 levels normally decline sharply^48,77^. This stabilization enables CDC20 to localize to the midbody in infected cells, where it colocalizes with E4orf4. Together, these findings suggest that AdV reprograms the temporal control of CDC20 to repurpose APC/C activity during late mitosis.

A key mechanistic insight from our study is that E4orf4 does not appear to act as an APC/C substrate but instead functions as an adaptor that redirects APC/C-CDC20 activity toward specific cytokinetic targets. We show that E4orf4 itself is not detectably ubiquitylated during infection or when expressed alone, although it can be stabilized by proteasome inhibitors, most likely through indirect effects. Instead, E4orf4 promotes robust hyperubiquitination of Aurora B kinase (AurB), a core component of the chromosomal passenger complex (CPC). AurB plays a central role in coordinating abscission timing through the abscission checkpoint, ensuring that cytokinesis is delayed until chromatin clearance from the midbody and cytoskeletal remodeling are complete^30,66^. While AurB is a known APC/C substrate, it is typically targeted by APC/C-CDH1 during late mitosis and G1. Whereas earlier results had shown an early activation of CDC20 by AdV E4orf4^50^, our data suggest that AdV infection enhances and recruits CDC20 levels via E4orf4 to the midbody, enabling premature and excessive APC/C-CDC20-dependent ubiquitylation of AurB at the midbody. Pharmacological inhibition of APC/C activity using Apcin or proTAME suppressed AurB hyperubiquitination and reduced cytokinetic regression, with proTAME strongly blocking E4orf4-CDC20 interaction. These results indicate that an active APC/C complex is required both for AurB modification and for E4orf4 function in promoting regression, underscoring APC/C-CDC20 as a central player in this viral manipulation. Our data further suggest that E4orf4 engages CDC20 in proximity to the C-terminal IR-tail, although the precise binding interface and critical residues remain to be defined. Elucidating this interaction will be important not only for understanding AdV-mediated cell cycle perturbation but also for identifying other viral or pathogenic proteins that may exploit APC/C modulation through a similar mechanism. Targeting this regulatory region might also provide novel ways to perturb cancer cell proliferation.

Live-cell imaging of endogenously tagged AurB reveals that, in AdV-infected cells, AurB initially localizes to the cleavage furrow but fails to maintain a stable midbody-associated pool. Instead, AurB signal became diffuse and was prematurely lost from the midbody, coinciding with peak E4orf4 accumulation, increased ubiquitin residence, and subsequent cleavage furrow regression. This behavior contrasts sharply with non-infected cells, where AurB remained stably associated with the midbody until abscission was complete. Previous studies had shown that premature inactivation or removal of AurB can override the abscission checkpoint and lead to cytokinetic failure or regression^30,31^. Our data extend these findings by demonstrating that a virus can actively exploit this vulnerability by driving AurB hyperubiquitination and extraction. The accumulation of ubiquitin at the midbody immediately prior to regression strongly supports a model in which AurB is removed in a highly ubiquitylated state, destabilizing midbody architecture and triggering daughter cell collapse.

Together, our findings support a model in which AdV hijacks the APC/C-CDC20 machinery via E4orf4 to perturb late cytokinesis. Whereas recent report has suggested that DNA viruses can inhibit APC/C activity outright to induce sustained mitotic arrest^78^, AdV appears to have evolved a distinct strategy that repurposes APC/C-CDC20 function to actively drive infected cells through abortive cytokinesis and into a polyploid state associated with enhanced viral gene expression. By stabilizing CDC20, forming an E4orf4-CDC20 complex at the midbody, and redirecting APC/C activity toward AurB, AdV induces premature midbody destabilization, cleavage furrow regression, and polyploid cell formation. Rather than representing a dead-end or cytotoxic outcome, this failure of cytokinesis generates large, viable cells that efficiently support viral gene expression. More broadly, this work identifies cytokinesis as an underappreciated target of viral manipulation and reveals APC/C-CDC20-AurB signaling as a vulnerable regulatory axis that can be exploited by pathogens. Given the central role of these pathways in maintaining genomic stability, their viral subversion may also have longer-term consequences for tissue homeostasis. Future studies will be required to define the CDC20 interface engaged by E4orf4, determine whether similar APC/C-rewiring strategies are employed by other viruses, and assess whether this pathway could be therapeutically targeted in virus-induced or cancer-associated cell cycle dysregulation.

### Limitations

Additional APC/C-CDC20 targets in the infected midbodies can drive the regression events along with AurB. In addition, other E4orf4-driven regulators of cytokinesis may contribute to AdV-induced cleavage furrow regression beyond the APC/C-CDC20 axis identified here. Indeed, our interactome analysis reveals several candidate E4orf4-interacting proteins with established roles in cytokinetic control; for example, PKP4, which has previously been implicated in cleavage furrow regression^79^, suggesting that its perturbation may account for the residual regression in CDC20 knockdown or inhibited cell. Finally, important caveats must be considered when interpreting the oncogenic implications of these findings, as polyploidy observed in *in vitro* culture or tissue models does not necessarily translate to tumorigenesis *in vivo*.

## Material and Methods

### Cell culture and transfection

Unless indicated, all cells were cultured in Dulbecco’s Modified Eagle’s Medium (DMEM) sup-plemented with 10% fetal calf serum (FCS), 100U/ml penicillin, 10µg/ml streptomycin, 1mM sodium pyruvate and non-essential amino acids. A549 cells with stable expression of indicated constructs (Key resources table) were created by lentiviral transduction. For lentivirus production, HEK293T cells were transfected with the lentiviral vector, along with expression constructs pMCV-Gag-Pol and pMD2-VSV-G (kind gifts from Dr Didier Trono, EPFL, Lausanne, Switzerland) using polyethyleneimine (PEI). Forty-eight hours after transfection, cell supernatant was passed thought 0.45µm pore-size filter and A549 cells were inoculated with filtered supernatant. After 24h, antibiotics were added: neomycin (500µg/ml), Blasticidin (10µg/ml), or Puromycin (1µg/ml) to select for indicated selections.

For siRNA mediated mRNA knockdown, pools of oligonucleotides (30nM) from Dharmacon (Horizon Discovery) were transfected into A549 cells using Lipofectamine RNAiMAX transfection reagent (Invitrogen) according to the manufacturer’s protocol. Two days post-transfection, cells were infected with indicated AdV strain at indicated MOIs, followed by timelapse imaging with IXM-C, Zeiss Celldiscover 7, Agilent Lionheart FX, or Olympus ScanR microscopes with 15min time resolution between frames, followed by quantification of the normal and regressing cells as described under “***Quantification of timelapse imaging***”.

### Human Bronchial Airway Basal Cell Culture and Adenoviral Infection

Human primary bronchial airway (HBA) basal cells were collected from an adult healthy donor (#AB0839, Epithelix) and expanded in PneumaCult Ex-Plus Medium (STEMCELL Technologies) at 37°C; 5% CO2. For live-cell time-lapse imaging, 20,000 cells were seeded per well in 96-well plates (CellVis). Two days post-seeding, cells were infected with AdV-C5-GFP-E4orf4 at the indicated multiplicities of infection (MOI). Time-lapse imaging was initiated 4 hours post-infection (hpi) and continued until 48 hpi using an ImageXpress system equipped with a 20X objective. For fixed-cell experiments, 20,000 cells were seeded per well in 96-well plates (CellVis) and infected with AdV-C5-GFP-E4orf4 at the indicated MOI. At 48 hpi, cells were fixed with paraformaldehyde (PFA), stained with Alexa Fluor™ 770–conjugated wheat germ agglutinin (WGA; 1:250 dilution in PBS) and Hoechst 33342 (1:2000), and imaged using the widefield mode of the ImageXpress system with a 60X objective. For DNA content analysis, cells were processed identically and imaged using the confocal mode of the ImageXpress system with a 20X objective and a 50 µm pinhole.

### Plasmids

The CDC20-mScarlet expression construct was generated from a human cDNA library generated from A549 cells as template using primers and inserted into the pLVX-mScarlet-N-term expression vector. Stable expression of cDNA in target cells was achieved using lentiviral transduction as described above^39^. Other variants of CDC20 expressing constructs including the bacterial expression were generated using this construct as a template. For the cloning of E4-promoter and E4 region, a 3025bp sequence from the C-terminal region of AdV-C5-GFP-E4orf4 was amplified via PCR and inserted in pLVX-IRES-Puro expression construct, replacing the constitutive CMV promoter. For validation, the construct was transfected in HeLa cells with or without exogenous E1A co-transfection, and expression of GFP-E4orf4 was confirmed via microscopy and western blotting, which occurred only in the presence of E1A expression. E4orf4 deletion construct in the same background was created by removing 345bp E4orf4 open-reading frame using site-directed mutagenesis, leaving the GFP expression intact. The conformation of ΔE4orf4 and WT-E4 region constructs in HEK293T cells were confirmed using microscopy and western blotting (**Fig. S8B**). Bacterial expression construct for E4orf4, CDC20, and E1A was performed using restriction cloning. Briefly, E4orf4 open-reading frame from template DNA extracted from purified AdV-C5_wt, 110-499 residues of CDC20 from human cDNA library, and full-length E1A-WT containing region was extracted from purified AdV-C5_wt genomic DNA and inserted in pET28a-10x-His plasmid construct using NheI-XhoI restriction enzyme pairs. All mammalian expression constructs of AurB and CDC20 were cloned via restriction cloning by extracting the open-reading frames from a human cDNA library. All resulting vectors were confirmed using Sanger sequencing.

### MMEJ-based knock-in of mScarlet at the C-term of Aurora B genomic locus

MMEJ-assisted knock-in for AurB was performed as described in Sakuma et al^63^. Plasmids pCRIS-PITChv2-FBL (#63672), pX330A-1×2 (#58766), and pX3301S-2-PITCh (#63670) were bought from Addgene. mScarlet ORF was inserted in place of EGFP in pCRIS-PITChv2-FBL using HiFi-DNA assembly. Left (5’-CTGCCCTTCAATCTGTCGCC-3’) and right (5’-TGATGGTCCCTGTCATTCAC-3’) microhomologies were inserted at the left and right arms of mScarlet-T2A-Puro cassette in modified pCRIS-PITChv2-FBL plasmid using HiFi-DNA assembly, hence generating pCRIS-PITChv2-AurB (Fig S9A). Guide RNA at the C-term of AurB (CCCTTCAATCTGTCGCCTGA) was inserted in pX330A-1×2 followed by transfer of U6-promoter-gRNA-scaffold to pX3301S-2-PITCh. HEK293T cells were seeded on 10-cm dish and were transfected with 1.2µg of pX330A0-1×2-gAurB-gPITCH and 0.6µg of pCRIS-PITChv2-AurB using Lipofectamine 2000. Three days post transfection, puromycin (2µg/ml) were added to the cells until resistance clones were observed. Clonal cells were picked from polyclonal resistant population via limiting dilution and AurB-mScarlet insertion was confirmed by genotyping and immunofluorescent imaging (Fig S9B).

### Correlative light and electron microscopy (CLEM)

CLEM was performed as described before^80^. In brief, A549 cells stably expressing AurB-mScarlet were grown on 35 mm-diameter dishes with gridded coverslip (Matek Corporation) and infected with AdV5-GFP-E4orf4 (moi5). Twenty-four hours later, cells were fixed with 4% paraformaldehyde and 0.2% glutaraldehyde (GA) in PBS for 30 min at room temperature, followed by washing with 150 mM glycine in PBS, once with PBS, stained with LipidToxTM Deep Red Neutral Lipid Stain (Invitrogen) and analysed using a spinning disk confocal microscope. Fluorescence images were acquired using a 100x objective with optical sections of 0.2mm. After imaging, cells were fixed again with 2.5% GA, 2% sucrose in 50 mM cacodylate buffer (cacodylate acid sodium trihydrate in distilled water, pH 7.2-7.4), supplemented with 50 mM KCl, 2.6 mM MgCl2 and 2.6 mM CaCl2 for at least 30 min on ice. Further processing for electron microscopy is described below. For correlation of fluorescent fiducial markers with EM ultrastructure markers, the ‘Landmark correspondence’ plugin in the ImageJ software package was used.

### Electron microscopy

Cells were fixed as described above, washed 5 times with 50 mM cacodylate buffer, incubated with 2% OsO4 in 25mM cacodylate buffer for 40min on ice, washed 3 times with water and incubated in 0.5% uranyl acetate in water overnight at 4°C. Samples were rinsed 3 times with water, dehydrated in a graded ethanol series (from 40 to 100%) at room temperature, followed by embedding in Epon 812 resin (Electron Microscopy Sciences). After polymerisation for 2 days at 60°C, cells of interest were located with the help of the imprint of the grid pattern on the resin block and either thin sections (70nm) for TEM or thicker sections (250nm) suitable for electron tomography were generated using a Leica EM UC6 ultramicrotome (Leica Microsystems). Sections were mounted on a slot grid and further counterstained using 3% uranyl acetate in 70% methanol for 5 min and lead citrate (Reynold’s) for 2 minutes. Images were taken with a JEM1400 microscope (JEOL GmbH) with a 4k TemCam-F416 camera (Tietz Video and Image Processing Systems GmbH). Images taken at 4,000X and 10,000X were stitched and correlated or quantified as described in respective sections.

### High-content and confocal Light microscopy

Confocal light microscopy was performed as described previously^80^. Briefly, cells seeded onto coverslips, 96-well dishes (CellVis), or 8-well dishes (Ibidi) were fixed using 4% paraformaldehyde, permeabilised with 0.1% Triton X-100 in PBS, blocked with 1% milk in PBS and incubated with primary antibody at room temperature for 60 min. Samples were washed with PBS three times and incubated with Alexa-dye conjugated secondary antibodies at room temperature for 30 min. After washing with PBS three times, PBS was added in 96-well or 8-well imaging plates and coverslips were mounted on imaging slides using Fluoromount-G (SouthernBiotech). Samples were imaged with the following microscopes: Agilent Lionheart FX (Fig. 4A-D, S3, S6, S7A) Zeiss CellDiscoverer 7 (Fig. 3F, 7D, S8B, S1B, S11, S2B), Zeiss spinning disk confocal (Fig. 1A, 1B, 4E, 6E, 6F, 7A, S9B, S10A-B), or Molecular device IXM-C (Fig. 1C-F, 2A, 2C, S1C, S2A, S8A).

### Quantification of time-lapse imaging

Time-lapse phase-contrast movies were quantified using a semi-automatic workflow combining manual annotation in Fiji and downstream data processing in KNIME. Individual mitotic events were manually annotated in Fiji using the “multi-point tool” by placing markers at the position and frame corresponding to cell division. Each division was classified by visual inspection as either regressing or normal cytokinesis. Marker identities were encoded numerically, with marker value 0 assigned to regressing divisions and 1 to normal divisions, and visualized with distinct colors (yellow or red, respectively). Using the Fiji Measure function, annotations were exported as a CSV file containing the marker identity (0 or 1) together with the corresponding time frame information. This CSV file was subsequently imported into KNIME, where annotations from each time-lapse were merged and processed to extract the total number of normal and regressing divisions per movie, as well as the timing of individual division events. This workflow enabled reproducible quantification of division fate and temporal dynamics across multiple time-lapse datasets.

### Correlative live and fixed imaging

To find if regressing cells accumulate more viral proteins than non-regressing cells during AdV infection, we performed live time-lapse imaging of AdV-infected cells lines (HeLa FUCCI, HeLa H2B-mCherry, or HeLa WT) were grown on 96-well imaging plates (Greiner Biosciences and CellVis glass bottom plates), infected with indicated AdV strain, followed by timelapse imaging with high-content microscope. After this, the cells were fixed, protein VI immunostaining was performed, nuclei were visualized with DAPI, and reimaging was performed with high-content microscope using the same settings and position of different wells as the timelapse. Transmission light channel with cellular morphology was used as the channel to align the last timepoint of time-lapse and fixed immunostained image. The alignment was done using CellProfiler. Post-alignment, the annotated cell number with cell fate as described in “Quantification of timelapse imaging” was correlated to the same cell and its protein VI mean intensity. The resulting graph was plotted using GraphPad Prism.

### Cell cycle stage determination from FUCCI cells

Cell cycle stage determination from fixed cells or individual timepoints of time-lapse imaging was done as described previously^16,81^. In brief, AdV-infected HeLa-FUCCI or HeLa-FUCCI-CA cells were fixed with formaldehyde post time-laspe and nuclei were visualized with DAPI. Nuclear stain of DAPI was used to segment cells using Cell Profiler. Quantification of cells in different cell cycle stages was performed using Cell Profiler Analyst software and Machine Learning Algorithms. FUCCI plasmid in HeLa cells expresses Kusabira orange fused Cdt1 gene (mKO-Cdt1) which indicates G1 stage, and Azami Green-labeled Geminin (mAGhGem), which indicates S/G2/M phase of cell cycle. A small gap in fluorescence change from red to green indicates a G1/S transition phase, already indicating an onset of S phase 12. Using these set of rules, the software differentiated between G1, G1/S and S/G2/M cells. Following this, the analysis was re-examined and errors were manually corrected and software was re-trained. After several iterations, the modified training set was used for the entire data set.

### GFP-E4orf4 intensity quantification in timelapse imaging

HeLa H2B-mCherry cells were infected with AdV-C5-GFP-E4orf4 and subjected to time-lapse imaging using a Molecular Devices ImageXpress Micro XL (IXM-XL) system, acquiring transmitted light, GFP, and TRITC channels at defined intervals. Regressing cell divisions were manually identified as described in the section “**Quantification of time-lapse imaging**” For quantitative analysis, transmitted light, H2B-mCherry, and GFP channels were merged and exported as RGB image stacks. Individual cells were cropped into fixed-size square regions (200 × 200 pixels) compatible with downstream analysis and split into individual time points with unique identifiers. Cell outlines were segmented from the transmitted light channel using Cellpose^82^, followed by manual verification and correction when necessary. Mean GFP-E4orf4 and H2B-mCherry intensities were measured at the single-cell level using CellProfiler^83^. Cytoplasmic and nuclear measurements were merged using KNIME^84^, and GFP-E4orf4 intensities were normalized to H2B-mCherry to account for cell-to-cell variability. For five randomly selected regressing cells exhibiting distinct division and regression timings, KNIME was used to determine the peak GFP-E4orf4 intensity, align intensity profiles to this peak, and map the timing of cleavage furrow regression relative to the normalized signal. Final plots were generated using JMP version 18.

### E4orf4 nuclear cytoplasmic fractionation

HeLa ATCC cells were seeded at a density of 3e6 cells per 10cm dish. They were infected next day with AdV-C5-GFP-E4orf4 (moi 5; 200 µL + 1.3ml complete DMEM) or mock. 36h post infection when GFP signal (indicating infected cells) was observed in almost all cells were lysed in 100µl NP40 lysis buffer (containing 0.1%-NP40). Later, lysates were incubated on ice for 30 mins and centrifuged at 16k g for 10mins to collect supernatant as cytoplasmic fraction and stored at −20°C. Washed the pellet (nuclei) with 100 µL of NP40 Lysis buffer and centrifuged at 16k g for 10 mins (performed only for technical replicate 1). Supernatant was removed and collected as wash fraction in a fresh tube. The nuclear pellet from both replicates was resuspended in RIPA buffer (50 µL) for 15mins. Vortexed briefly 2-3 times in between. After 15 mins centrifuged at max speed for 10 mins to extract nuclear fraction, sup collected in a fresh tube and stored at −20. The resulting pellet was also stored. Cytosolic and nuclear extracts were quantified with DC (detergent compatible) protein assay buffers from Bio-Rad (#5000111) and 40µg of total protein from each was loaded on SDS-PAGE together with nuclear and cytoplasmic controls to check for purity of cytoplasmic and nuclear extracts. The cytoplasmic extracts were utilized for anti-GFP immunoprecipitation and processed for liquid chromatography-tandem mass spectrometry (LC-MS/MS) analysis at DKFZ, Heidelberg.

### GFP-trap immunoprecipitation of E4orf4

For LC-MS/MS analysis of cytoplasmic E4orf4, 3 million HeLa cells were cultured in 10-cm dishes and mock-infected or Adenovirus-GFP–E4orf4 infected in parallel. 30 h post-infection cells were harvested under cold conditions, rinsed with phosphate-buffered saline, and lysed using a non-ionic detergent lysis buffer supplemented with protease inhibitors to obtain clarified cytoplasmic extracts as described before. A portion of each lysate was reserved as input, and the remaining material was subjected to GFP immunoprecipitation by incubating the extracts with an anti-GFP antibody followed by capture on Protein A resin overnight at 4°C on end-to-end rotator. After removal of nonspecifically bound material through sequential washes, bound proteins were eluted under denaturing conditions. Input and immunoprecipitated samples were subsequently processed for SDS–PAGE to evaluate sample quality of cytoplasmic and nuclear extracts in mock and infected conditions. 2x Laemmli dissolved and boiled (95°C) cytoplasmic extracts of both mock and infected samples were processed for liquid chromatography-tandem mass spectrometry (LC-MS/MS) analysis at DKFZ, Heidelberg (described below in “*Mass spectrometry-based analysis*”).

For other immunoprecipitations, HeLa-ATCC, A549 or 293T cells expressing tagged proteins indicated in different figures were seeded on 10-cm dishes. Twenty-four hours later, cells were infected with AdV-C5-GFP-E4orf4 (moi 5) or transfected with pE4-Ad-E4 or pE4-Ad-E4-E4orf4-KO plasmids at 37 °C, 5% CO2 for 24 h. Following this, the medium was aspirated and washed with ice-cold PBS several times. Cells were scraped in 600 μl of DMEM and collected in 1.5-ml tubes. Cells were pelleted and resuspended in 600 μl of ice-cold IP lysis buffer (Tris-HCl, pH 8.5, NaCl 150 mM, MgCl2 1 mM, EDTA 1 mM, Nonidet P-40 0.5%, and protease inhibitor cocktail) under agitation at 4 °C for 20 min. The cells were centrifuged at 16,000×g at 4 °C for 10 min, and the supernatant was collected. Fifty microliters of the cell lysates were kept for an input control sample and the rest was precleared with protein-A Dyna beads (ThermoFisher, #`10001D`). Precleared lysates were incubated with anti-HA (Sigma, F7425) overnight at 4 °C. The next day, 30 μl of Protein-A Dynabeads were added per sample and incubated at 4 °C for 1 h with agitation. Beads were spun down, the supernatant removed, and beads washed in IP lysis buffer several times. After this, the beads were mixed with 2×SDS lysis buffer (10% w/v), boiled at 95 °C for 5 min, and the supernatant was collected. Samples were resolved with polyacrylamide gel electrophoresis after addition of DTT (50 mM) for immunoblotting.

### Mass spectrometry-based analysis

Immunoprecipitation (IP) eluates (20 µl) were processed for proteomic analysis using an automated single-pot solid-phase–enhanced sample preparation (autoSP3) workflow implemented on an AssayMAP Bravo liquid handling system (Agilent Technologies), including tryptic digestion, according to Müller et al^85^. Dried peptide samples were reconstituted in 97.4% water, 2.5% hexafluoro-2-propanol, and 0.1% trifluoroacetic acid, and an amount corresponding to 20 µl of the original IP eluate was injected for LC–MS/MS analysis. Peptides were separated on either an Ultimate 3000 or a Vanquish Neo UPLC system (Thermo Fisher Scientific) directly coupled to an Orbitrap Exploris 480 mass spectrometer (Thermo Fisher Scientific), using a total run time of 60 min per sample. Online desalting was performed on an Acclaim PepMap300 C18 trapping cartridge (5 µm, 300 Å; Thermo Fisher Scientific) using 0.05% TFA in water at 30 µl/min for 3 min (Ultimate 3000; 30 µl loading) or with a 60 µl loading volume (Vanquish Neo). Peptide separation was achieved at 300 nl/min on a nanoEase MZ Peptide analytical column (300 Å, 1.7 µm, 75 µm × 200 mm; Waters) using solvent A (0.1% formic acid in water) and solvent B (0.1% formic acid in acetonitrile). For Ultimate 3000 runs, solvent B was linearly increased from 5% to 30% over 46 min, ramped to 78%, reduced to 2% after 2 min, and followed by a 5 min equilibration. For Vanquish Neo runs, solvent B was ramped from 5% to 30% over 45 min, increased to 80%, reduced to 2% after 4 min, and equilibrated for three column volumes. Mass spectrometric data were acquired in data-dependent acquisition (DDA) mode with a full MS scan at 60,000 resolution (m/z 380–1400, 500% AGC target, 100 ms maximum injection time), followed by up to 2 s of MS/MS acquisition. Precursors were isolated with a 1.2 m/z window and fragmented using 26% normalized collision energy. Fragment spectra were acquired at 15,000 resolution (100% AGC target, 150 ms maximum injection time). Unassigned and singly charged precursors were excluded, and dynamic exclusion was set to 10 s. Each sample was followed by a 20 min wash run to minimize carry-over. Instrument performance was monitored throughout data acquisition by regular injections (approximately every 48 h) of a standard sample and evaluated using an in-house Shiny-based quality control application.

For data analysis, raw data were processed using MaxQuant (version 2.1.4.0)^86^ with default parameters unless otherwise stated. Searches were performed against organism-specific UniProt databases comprising the human reference proteome (20,597 entries, one protein per gene; downloaded 9 February 2024) and Human adenovirus C serotype 5 (488 entries). Match-between-runs was enabled to transfer peptide identifications based on accurate mass and retention time, with fraction settings restricting matching to within biological replicates. Label-free quantification (LFQ) was enabled using default settings, and intensity-based absolute quantification (iBAQ) values were calculated^87^.

### Western blotting

Cells were cultured in 24-well plates and either infected or transfected with the indicated constructs. Whole-cell lysates were prepared using lysis buffer (0.2 ml containing 200 mM Tris pH 8.8, 20% glycerol, 5 mM EDTA, 50 mM DTT, 5% SDS, 0.02% bromophenol blue), followed by boiling at 95 °C for 5 min and mechanical shearing through G21 needles (Sterican). Proteins were separated on 10–15% SDS–polyacrylamide gels depending on molecular weight, transferred onto polyvinylidene difluoride (PVDF) membranes (Amersham), and blocked in 5% milk. Immunoblotting was performed using antibodies against E1A (M58), HA, GFP, Flag, GAPDH, and β-tubulin.

For biochemical analyses, cells were rinsed once with ice-cold PBS and collected by scraping into protein lysis buffer (50 mM Tris-HCl pH 7.4, 150 mM NaCl, 1% Triton X-100, 60 mM β-glycerophosphate, 15 mM 4-nitrophenylphosphate, 1 mM Na_₃_VO_₄_, 1 mM NaF in dH_₂_O), supplemented with a protease inhibitor cocktail (Roche). Lysates were incubated on ice for 30 min and clarified by centrifugation at 20,800 × g for 15 min. Supernatants were transferred to fresh tubes and stored at −80 °C. Protein concentrations were determined using Bio-Rad protein assay dye. Equal amounts of protein (100 µg) were combined with 6× sample buffer (375 mM Tris-HCl pH 6.8, 50% glycerol, 9% SDS, 9% β-mercaptoethanol, 0.03% bromophenol blue) and resolved on 8% SDS–polyacrylamide gels using 1× TGS running buffer at 80–120 V.

Proteins were electrotransferred onto methanol-activated PVDF membranes using transfer buffer (2.5 mM Tris base, 15 mM glycine pH 8.3, 20% methanol) at 360 mA for 2 h. Membranes were blocked for 1 h at room temperature in TBS-T containing either 5% BSA or 5% non-fat dry milk. Primary antibodies (listed in the Key Resources table) were diluted in 5% milk-TBS-T (0.1% Tween-20) and incubated overnight at 4 °C. After three washes in TBS-T, membranes were incubated with HRP-conjugated secondary antibodies for 1 h at room temperature, washed again, and signals were visualized using Western Lightning chemiluminescence substrate (PerkinElmer) and imaged on INTAS or Bio-Rad systems. Band intensities were quantified using LabImage1D (INTAS) or Image Lab (Bio-Rad) software and normalized to the corresponding loading controls. For each biological replicate, values were additionally normalized to the total signal per membrane to correct for inter-replicate variability and are presented as scaled arbitrary units (AU).

### RNAi knockdown of CDC20 and timelapse imaging

siRNA-mediated gene silencing was performed using the Cherry Pick Library (Dharmacon) in a 96-well format with A549-ACE2 reporter cells. Library siRNAs were diluted to generate working stocks (2µM) before transfection. For each siRNA condition, a lipid-based transfection mixture was prepared using Opti-MEM, Lipofectamine RNAiMAX, and siRNA at a final concentration of 30nM. Following a short incubation of 20 minutes to allow complex formation, A549-ACE2 cells were harvested, counted, and adjusted to a density suitable for seeding in 96-well plates. Transfection complexes were dispensed into duplicate plates, after which cells were added directly to each well and maintained under standard culture conditions for 48 h to allow gene silencing. After this period, one plate was fixed with 4% paraformaldehyde for analysing nuclear and cellular morphology using Hoechst and wheat germ agglutinin staining, whereas the second plate was infected with wild-type Adenovirus for subsequent live-cell imaging. Live imaging of infected cells was conducted over a two-day period with Zeiss CellDiscoverer 7, and regression frequencies were quantified as described in section “***Quantification of timelapse***”.

### RNA isolation and quantitative PCR (qPCR)

As directed by the manufacturer, total RNA was extracted using the NucleoSpin RNA kit (Macherey-Nagel). Thermo Fisher Scientific’s High-Capacity cDNA Reverse Transcription kit was used to create complementary DNA (cDNA) from whole RNA. Using iTaq Universal SYBR Green Supermix (Bio-Rad), cDNA was subjected to qPCR analysis. Primers specific to GAPDH (forward: 5′-GAAGGTGAAGGTCGGAGTC-3′; reverse: 5′-GAAGGTGGTGATGGGATTTC-3′) and CDC20 (forward: 5′-CGGAAGACCTGCCGTTACATTC-3′; reverse: 5′-CAGAGCTTGCACTCCACAGGTA-3′) were used. Melt-curve analysis was used to confirm the amplification specificity of the reactions. Relative transcript abundance was normalized to GAPDH. The comparative Ct (2^−ΔΔCt) method was used to calculate the relative mRNA expression levels of CDC20. By comparing CDC20 expression in knockdown samples to non-targeting siRNA control cells, which were adjusted to 1, knockdown efficiency was calculated.

### Ni-NTA-based immunoprecipitation for His-tagged protein purification

A Ni–NTA pull-down assay was performed to isolate His-tagged Ubiquitylated proteins expressed in HEK293T cells. 5e6 cells were seeded per condition in a 10cm dish and transfected with the indicated plasmids. 24h post transfection cells were lysed by several freeze-thawing under denaturing conditions in Urea Buffer 1, which consisted of 8 M urea, 100 mM sodium phosphate (NaH_₂_PO_₄_), 10 mM Tris-Cl at pH 8.0, and 25 mM imidazole, supplemented with protease inhibitor cocktail. Lysates were clarified and incubated with 30µl (per sample) Ni–NTA dyna beads (ThermoFisher, #88832) to allow binding of His-tagged proteins at room temperature. After 4h of binding at end-to-end rotator, the resin was washed thrice with Urea Buffer 2 containing 8 M urea, 100 mM NaH_₂_PO_₄_, 10 mM Tris-Cl adjusted to pH 6.3, and 30 mM imidazole, providing stringent removal of nonspecifically associated proteins. Bound material was eluted in 50µl Urea Buffer 3, composed of 8 M urea, 100 mM NaH_₂_PO_₄_, 10 mM Tris-Cl at pH 4.0, and 300 mM imidazole. 2×SDS lysis buffer was added to eluted and input samples, boiled at 95°C followed by subsequently resolving by SDS–PAGE and analyzing by immunoblotting with the indicated antibodies.

### Cell synchronization, infection, and drug treatments

Cells were seeded in a glass bottom 96-well plate at a density of 6000 cells per well and synchronized using a double thymidine block. Briefly, cells were treated with 2 mM thymidine (Sigma-Aldrich; T9250) for 18 h, released into fresh medium for 9 h, and subjected to a second thymidine block for an additional 18 h to arrest cells in S phase. Cells were infected with GFP-E4ORF4 or Ad2WT virus at the indicated Infectious amount approximately 4 h before the final release. Thymidine was removed at 18 h post-block, coinciding with drug addition. Where indicated, cells were treated with Apcin (Sigma-Aldrich, SML1503; 50µM) or ProTAME ((Sigma-Aldrich, SML3977; 12 µM) at the time of thymidine removal. Under these conditions, cells progressed synchronously through early mitosis, reaching mitosis approximately 9 h after release. Non-infected controls and vehicle-treated samples were included in parallel. 24 hours post-infection, cells were fixed with 16% paraformaldehyde (PFA) on top for 15 minutes at room temperature in the dark. After fixation, permeabilisation was performed using 0.1% Triton X-100 in PBS for 5 minutes at room temperature, then incubated with CDC20 antibody (Labforce; sc-1316; 1:200 in 1% skim milk in PBST) incubated overnight at 4°C overnight in rocker. Following primary antibody incubation, cells were washed three times with PBS and incubated Goat anti-Mouse IgG (H+L) Cross-Adsorbed Secondary Antibody, Alexa Fluor 750 (1:1000 dilution in PBS + 1% skim milk) for 1 hour at room temperature on a rocker in the dark. Nuclei were stained with Hoechst (1:1000 dilution from a 1 mg/ml stock) followed by a final PBS wash.

### CDC20 protein estimation in G2/M synchronized cells

A549 cells were seeded at a density of 4 × 10LJ cells per 10-cm dish and synchronized by thymidine arrest using 2.5 mM thymidine for 24 h. Cells were released into thymidine-free medium and simultaneously infected with Ad2WT virus at the indicated Infectious amount. Two hours post-release, 10 µM S-trityl-L-cysteine (STLC; Sigma-Aldrich 164739) were added to arrest cells in spindle assembly checkpoint. Cells were maintained under STLC treatment for 16 h, after which lysates were collected. Cells were pelleted by centrifugation and lysed in 300 µl IP lysis buffer and 1x PIC. DNase I (Sigma-Aldrich, 10104159001; 0.1 mg/mL) added to the lysate. Protein concentrations were estimated by Bradford assay. Equal concentration of cell lysates were loaded and separated by SDS–PAGE and transferred onto a 0.22-µm nitrocellulose membrane. After transfer, the membrane was blocked with 5% skim milk in TBST (TBS containing 0.1% Tween-20) for 1 h at room temperature. It was then incubated overnight at 4 °C with an anti-CDC20 primary antibody CDC20 antibody (Labforce; sc-1316; 1:500 in 5% skim milk in PBST) and Adenovirus-5 E1A (M58) (SantaCruz; sc-58658; 1:1000 in 5% skim milk in PBST). Following washes with TBST, the membrane was incubated for 1 h at room temperature with a Goat Anti-Mouse IgG (H+L), Peroxidase Conjugated secondary antibody (Fisher, 10158113; diluted 1:2000 in 5% BSA/TBST. After three additional TBST washes, signals were developed using the Immobilion Forte Western HRP Substrate (Sigma-Aldrich WBLUF0100) according to the manufacturer’s protocol. Images were acquired using an ImageQuant LAS 500 system (GE Life Sciences, USA) and quantified with Bio-Rad Image software.

### Recombinant protein expression, purification, and *in vitro* immunoprecipitation

FLAG-E4ORF4, CDC20(110-499)-HA-GST, and E1A-wild-type genes, encoded in the pET28a plasmid were transformed in Escherichia coli Rosetta competent cells using the standard heat-shock protocol. Single colonies were used to inoculate 4 mL LB medium supplemented with kanamycin (50 µg/mL; Sigma-Aldrich, K1876-1G) and grown overnight at 37 °C with shaking. Expression cultures were initiated by diluting overnight cultures 1:50 into 100 mL LB containing kanamycin (50 µg/mL) and chloramphenicol (50 µg/mL; AxonLab, 10022944) and grown at 37 °C to an OD_₆₀₀_ (optical density at 600 nm) of 0.6.

For FLAG-E4ORF4 and CDC20(110-499)-HA-GST cultures were cooled on ice for 30 min and protein expression was induced with 0.5 mM isopropyl β-d-1-thiogalactopyranoside (IPTG; AxonLab, 10021074). Ethanol was added to a final concentration of 2% (v/v) to arrest further bacterial growth, and cultures were incubated overnight (at least 16 hours) at 16 °C with shaking. Cells were harvested by centrifugation at 4,700 rpm for 30 min at 4 °C, and pellets were stored at −80 °C prior to lysis. Cell pellets of FLAG-E4ORF4 were resuspended in 20 mL Buffer A (50 mM Tris–HCl pH 8.0, 500 mM NaCl, 10% glycerol) supplemented with lysozyme (0.5 mg/mL), Triton X-100 (0.1%), DNase I (Sigma-Aldrich, 10104159001; 0.1 mg/mL), DTT (AxonLab, 10021384; 0.2 mM), and 2x protease inhibitor cocktail (Sigma-Aldrich, 11836170001). CDC20(110-499)-HA-GST lysed in Buffer B (25 mM Tris–HCl pH 6.5, 250 mM NaCl, 1 mM MgCl2, 5% glycerol) supplemented with lysozyme (0.5 mg/mL), Triton X-100 (0.1%), DNase I (0.1 mg/mL), DTT (0.2 mM), and 1x protease inhibitor cocktail. For E1A Protein expression was induced by addition of IPTG to a final concentration of 1.5 mM, followed by incubation at 42 °C for 3 h. Cultures were cooled on ice for 30 min. Cells were harvested by centrifugation at 5,000 × g for 20 min at 4 °C, and pellets were stored at −80 °C. Cell pellets were resuspended in 20 mL Buffer C consisted of 50 mM HEPES (pH 8.0), 200 mM NaCl, 10% (v/v) glycerol, 1 mM EDTA (pH 8.0), and 10 mM MgCl_₂_. supplemented with lysozyme (0.2 mg/mL), DTT (0.1 mM), DNase I (15 µg/mL), and 1x protease inhibitor cocktail.

Chemical lysis for all the bacterial pellet was performed on ice for 1 hour with vortexing at every 10 minutes followed by two passes through a microfluidizer (LM20) at 18000 psi. Lysates were clarified by centrifugation at 12,000 rpm for 1 h at 4 °C. The soluble fraction was supplemented with imidazole to 20 mM and applied to Ni-NTA agarose resin equilibrated in the corresponding lysis Buffer specific for each protein. After binding by gravity flow, the flow-through was reapplied once. The resin was washed with Buffer A containing 20 mM imidazole, and bound proteins were eluted stepwise using Buffer A supplemented with 50-, 100-, 250-, or 500-mM imidazole. Eluted fractions, flow-through, washes, and resin-bound material were analysed and purity of the eluted protein was checked using SDS-PAGE.

### In vitro GST immunoprecipitation assays

20 µL of dry GST agarose beads (G0924-1ML; Sigma-Aldrich) were used for immobilization were equilibrated by washing three times with 0.5 mL of Buffer A (50 mM Tris–HCl, pH 8.0; 500 mM NaCl; 10% glycerol) with 0.1% Tx100 to remove storage preservatives and to equilibrate the beads in assay buffer. After equilibration, beads were incubated either alone as a GST beads control or with purified GST–CDC20 (residues 110–499; 450 µL at 13.4 ng/µL, corresponding to approximately 3.0 µg of protein per reaction) for 2h at 4 °C under continuous rotation. Following incubation, beads were collected using a magnetic stand, and the flow-through fractions were carefully retained for analysis. Beads were then washed three times with Buffer A containing 0.1% Triton X-100 and subsequently resuspended in 20 µL Buffer A. For interaction assays, 10 µL aliquots of the prepared beads were incubated in 250 µL Buffer A under the following conditions: (i) GST beads alone with 3.0 µg FLAG–E4ORF4, (ii) GST Beads + CDC20-GST with 3.0 µg FLAG–E4ORF4, and (iii) GST beads alone with 3.0 µg E1A and (iv) GST beads + CDC20-GST with 3.0 µg E1A protein as a negative control. These reactions were performed for 1 h at room temperature with gentle. After incubation, beads were collected using the magnetic stand, and the flow-through fractions were retained. Beads were washed three times, each time with 0.5 mL Buffer A containing 0.1% Triton X-100 to remove non-specifically bound proteins. Proteins bound to the beads were eluted by boiling in 2× Laemmli sample buffer at 95 °C for 10 min, followed by the addition of 1 mM dithiothreitol (DTT) to reduce disulfide bonds.

The flow through fractions along with the washes combined were subjected to TCA precipitation. one volume of TCA stock was added to four volumes of protein sample. Samples were incubated overnight at –20 °C to allow protein precipitation. Precipitated proteins were collected by centrifugation at 14,000 rpm for 5 min at 4 °C. The supernatant was carefully removed, leaving the whitish, fluffy protein pellet intact. If the pellet not visible at this stage, the supernatant was subjected to reprecipitation. Pellets were washed twice with 200 µL of cold acetone, centrifuging at 14,000 rpm for 5 min between washes, to remove residual TCA and salts. The final pellet was air-dried or placed in a 95 °C heat block for 5–10 min to remove acetone. Proteins were solubilized in 2× Laemmli buffer. Eluted samples and corresponding flow-through fractions were analysed by SDS–PAGE. Proteins were resolved on 12% SDS–PAGE gels, with samples loaded at a 3:1 ratio: the higher volume sample was used for Coomassie Brilliant Blue staining (Lubio Quick Blue Protein Stain,LJLU001000) to visualize total protein, and the lower volume sample was subjected to immunoblotting. Immunodetection was performed using anti-HA antibody (Thermo Fisher Scientific, PA1-985) overnight at 4 degrees 1:5000 in 5% skim milk in PBST to detect CDC20 and 1 h at room temperature for E1A. E1A Adenovirus-5 E1A (M58) (SantaCruz; sc-58658; 1:8000 in 5% skim milk in PBST) used for detection of E1A for 1 h at room temperature and anti-FLAG antibody for detection of FLAG–E4ORF4 (Sigma-Aldrich, F7425, 1:5000 in 5% skim milk in PBST 1 h at room temperature). Following washes with TBST, the membrane was incubated for 1 h at room temperature with a Goat Anti-Mouse IgG (H+L), Peroxidase Conjugated secondary antibody or Goat Anti-Rabbit IgG (H+L) (Fisher, 11859140; Fisher, 10158113; diluted 1:8000 in 5% skim milk in TBST). After three additional TBST washes, signals were developed using the Immobilion Forte Western HRP Substrate (Sigma-Aldrich WBLUF0100) according to the manufacturer’s protocol. Images were acquired using an ImageQuant LAS 500 system (GE Life Sciences, USA) and quantified with Bio-Rad Image software.

## Supporting information

Table S1: Proteins identified in GFP-E4orf4 proteomics

## Data availability

The generated proteomics data are deposited in PRIDE Proteomics database and accessible through accession number PXD073585.

## Acknowledgements

We thank all the members of microbiology and molecular medicine (MIMOL) at University of Geneva, the Molecular Virology research unit at Heidelberg University, and Molecular Life Sciences at University of Zurich for discussions and support. We gratefully acknowledge Prof. Dr. Dr. h.c. Ralf Bartenschlager for providing access to resources, including funding of personnel, access, and discussions with DKFZ proteomics facility, IDIP for imaging and EMCF for CLEM experiments. We thank Prof. Caroline Tapparel for discussions and providing access to primary cells. We thank Mr. Johan Geiser and Prof. Martina Valentini for discussions and assistance with protein expression and purifications. We thank Dr. Maarit Suomalainen for discussions. We acknowledge the support of the IDIP imaging core facility, headed by Dr. Vibor Laketa at the CIID, Heidelberg, Electron Microscopy Core Facility, Heidelberg, and the Bioimaging Core Facility at CMU, University of Geneva. We thank Prof. Daniel Gerlich, IMBA, Austria, for the provisions of cell lines, Prof. Phillip Branton and Dr. Paula Blanchette for the provision of viruses and Prof. Tamar Kleinberger, Israel Institute of Technology, Israel for the provision of cell lines.

This work was supported from grants from Kanton Zürich to U.F.G. and Olga Mayenfisch Stiftung (Zürich), Ernst Boninchi Foundation (Genève), and Faculty of Medicine and Department MIMOL (University of Geneva) startup funds to V.P.

## Author contributions

Conceived and conceptualized: V.P. Designed, performed, and analyzed experiments: S.S., D.S., Y.J., Va.Po., M.B., M.H., V.P. Expression, purification, and in vitro IP experiments: D.S. under the supervision of V.P. V.P. performed CLEM experiment and EM imaging and correlation, U.H. prepared thin sections for EM. M.S., D.H. performed mass spectrometry and analyzed results. Wrote the manuscript: V.P. with inputs from S.S., D.S. Discussions, editing, validation: M.S., M.B., U.F.G., V.P. Funding acquisition: U.F.G., V.P.

## Competing interests

The authors declare no competing interests.

## Notes

### Competing Interest Statement

The authors have declared no competing interest.

## References

1. Bagga, S., and Bouchard, M.J. (2014). Cell Cycle Regulation During Viral Infection. Cell Cycle Control 1170, 165–227. 10.1007/978-1-4939-0888-2_10.

2. Davoli, T., and de Lange, T. (2011). The causes and consequences of polyploidy in normal development and cancer. Annu. Rev. Cell Dev. Biol. 27, 585–610. 10.1146/annurev-cellbio-092910-154234.

3. Fujiwara, T., Bandi, M., Nitta, M., Ivanova, E.V., Bronson, R.T., and Pellman, D. (2005). Cytokinesis failure generating tetraploids promotes tumorigenesis in p53-null cells. Nature 437, 1043–1047. 10.1038/nature04217.

4. Patel, D., Incassati, A., Wang, N., and McCance, D.J. (2004). Human Papillomavirus Type 16 E6 and E7 Cause Polyploidy in Human Keratinocytes and Up-Regulation of G2-M-Phase Proteins. Cancer Res. 64, 1299–1306. 10.1158/0008-5472.CAN-03-2917.

5. Liu, M.-T., Chen, Y.-R., Chen, S.-C., Hu, C.-Y., Lin, C.-S., Chang, Y.-T., Wang, W.-B., and Chen, J.-Y. (2004). Epstein-Barr virus latent membrane protein 1 induces micronucleus formation, represses DNA repair and enhances sensitivity to DNA-damaging agents in human epithelial cells. Oncogene 23, 2531–2539. 10.1038/sj.onc.1207375.

6. Rakotomalala, L., Studach, L., Wang, W.-H., Gregori, G., Hullinger, R.L., and Andrisani, O. (2008). Hepatitis B virus X protein increases the Cdt1-to-geminin ratio inducing DNA re-replication and polyploidy. J. Biol. Chem. 283, 28729–28740. 10.1074/jbc.M802751200.

7. Pan, H., Zhou, F., and Gao, S.-J. (2004). Kaposi’s sarcoma-associated herpesvirus induction of chromosome instability in primary human endothelial cells. Cancer Res. 64, 4064–4068. 10.1158/0008-5472.CAN-04-0657.

8. White, M.K., Pagano, J.S., and Khalili, K. (2014). Viruses and Human Cancers: a Long Road of Discovery of Molecular Paradigms. Clin. Microbiol. Rev. 27, 463–481. 10.1128/cmr.00124-13.

9. Lion, T. (2019). Adenovirus persistence, reactivation, and clinical management. FEBS Lett. 593, 3571–3582. 10.1002/1873-3468.13576.

10. Sequeira, D.P., Suomalainen, M., Freitag, P.C., Plückthun, A., Klingenbrunner, M., Fischer, L., Hemmi, S., Münz, C., Volle, R., and Greber, U.F. (2025). Activated blood-derived human primary T cells support replication of HAdV C5 and virus transmission to polarized human primary epithelial cells. J. Virol. 0, e01825–24. 10.1128/jvi.01825-24.

11. Ben-Israel, H., and Kleinberger, T. (2002). Adenovirus and cell cycle control. Front. Biosci. J. Virtual Libr. 7, d1369–1395. 10.2741/ben.

12. Chellappan, S.P., Hiebert, S., Mudryj, M., Horowitz, J.M., and Nevins, J.R. (1991). The E2F transcription factor is a cellular target for the RB protein. Cell 65, 1053–1061. 10.1016/0092-8674(91)90557-F.

13. Bandara, L.R., and La Thangue, N.B. (1991). Adenovirus E1a prevents the retinoblastoma gene product from complexing with a cellular transcription factor. Nature 351, 494–497. 10.1038/351494a0.

14. Kafle, C.M., Anderson, A.Y., Prakash, A., Swedik, S., and Bridge, E. (2022). An Adenovirus early region 4 deletion mutant induces G2/M arrest via ATM activation and reduces expression of the mitotic marker phosphorylated (ser10) histone 3. Virology 565, 1–12. 10.1016/j.virol.2021.09.006.

15. Turner, R.L., Groitl, P., Dobner, T., and Ornelles, D.A. (2015). Adenovirus replaces mitotic checkpoint controls. J Virol 89, 5083–5096. 10.1128/JVI.00213-15.

16. Prasad, V., Suomalainen, M., Hemmi, S., and Greber, U.F. (2017). Cell Cycle-Dependent Kinase Cdk9 Is a Postexposure Drug Target against Human Adenoviruses. ACS Infect. Dis. 3, 398–405. 10.1021/acsinfecdis.7b00009.

17. Grand, R.J., Ibrahim, A.P., Taylor, A.M., Milner, A.E., Gregory, C.D., Gallimore, P.H., and Turnell, A.S. (1998). Human cells arrest in S phase in response to adenovirus 12 E1A. Virology 244, 330–342. 10.1006/viro.1998.9102.

18. Sriskandarajah, N., Blanchette, P., Kucharski, T.J., Teodoro, J.G., and Branton, P.E. (2015). Analysis by Live Imaging of Effects of the Adenovirus E4orf4 Protein on Passage through Mitosis of H1299 Tumor Cells. J. Virol. 89, 4685–4689. 10.1128/JVI.03437-14.

19. Graham, F.L., and Van Der Eb, A.J. (1973). Transformation of rat cells by DNA of human adenovirus 5. Virology 54, 536–539. 10.1016/0042-6822(73)90163-3.

20. Breuer, W., Trustaedt, M., and Hermanns, W. (2006). [Appearance of multinucleated giant cells in association with an adenovirus infection in two guinea pigs]. Berl. Munch. Tierarztl. Wochenschr. 119, 474–479.

21. Kleinberger, T. (2020). Biology of the adenovirus E4orf4 protein: from virus infection to cancer cell death. FEBS Lett. 594, 1891–1917. 10.1002/1873-3468.13704.

22. Shtrichman, R., Sharf, R., Barr, H., Dobner, T., and Kleinberger, T. (1999). Induction of apoptosis by adenovirus E4orf4 protein is specific to transformed cells and requires an interaction with protein phosphatase 2A. Proc. Natl. Acad. Sci. 96, 10080–10085. 10.1073/pnas.96.18.10080.

23. Mannervik, M., Fan, S., Ström, A.C., Helin, K., and Akusjärvi, G. (1999). Adenovirus E4 open reading frame 4-induced dephosphorylation inhibits E1A activation of the E2 promoter and E2F-1-mediated transactivation independently of the retinoblastoma tumor suppressor protein. Virology 256, 313–321. 10.1006/viro.1999.9663.

24. Cabon, L., Sriskandarajah, N., Mui, M.Z., Teodoro, J.G., Blanchette, P., and Branton, P.E. (2013). Adenovirus E4orf4 Protein-Induced Death of p53−/− H1299 Human Cancer Cells Follows a G1 Arrest of both Tetraploid and Diploid Cells due to a Failure To Initiate DNA Synthesis. J. Virol. 87, 13168–13178. 10.1128/JVI.01242-13.

25. Li, S., Brignole, C., Marcellus, R., Thirlwell, S., Binda, O., McQuoid, M.J., Ashby, D., Chan, H., Zhang, Z., Miron, M.-J., et al. (2009). The Adenovirus E4orf4 Protein Induces G2/M Arrest and Cell Death by Blocking Protein Phosphatase 2A Activity Regulated by the B55 Subunit. J. Virol. 83, 8340–8352. 10.1128/jvi.00711-09.

26. Mui, M.Z., Kucharski, M., Miron, M.-J., Hur, W.S., Berghuis, A.M., Blanchette, P., and Branton, P.E. (2013). Identification of the Adenovirus E4orf4 Protein Binding Site on the B55α and Cdc55 Regulatory Subunits of PP2A: Implications for PP2A Function, Tumor Cell Killing and Viral Replication. PLOS Pathog. 9, e1003742. 10.1371/journal.ppat.1003742.

27. Potapova, T., and Gorbsky, G.J. (2017). The Consequences of Chromosome Segregation Errors in Mitosis and Meiosis. Biology 6, 12. 10.3390/biology6010012.

28. Oldstone, M.B.A. (2006). Viral persistence: parameters, mechanisms and future predictions. Virology 344, 111–118. 10.1016/j.virol.2005.09.028.

29. Virgin, H.W., Wherry, E.J., and Ahmed, R. (2009). Redefining chronic viral infection. Cell 138, 30–50. 10.1016/j.cell.2009.06.036.

30. Steigemann, P., Wurzenberger, C., Schmitz, M.H.A., Held, M., Guizetti, J., Maar, S., and Gerlich, D.W. (2009). Aurora B-Mediated Abscission Checkpoint Protects against Tetraploidization. Cell 136, 473–484. 10.1016/j.cell.2008.12.020.

31. Carlton, J.G., Caballe, A., Agromayor, M., Kloc, M., and Martin-Serrano, J. (2012). ESCRT-III governs the Aurora B-mediated abscission checkpoint through CHMP4C. Science 336, 220–225. 10.1126/science.1217180.

32. Schaack, J., Guo, X., and Langer, S.J. (1996). Characterization of a replication-incompetent adenovirus type 5 mutant deleted for the preterminal protein gene. Proc. Natl. Acad. Sci. U. S. A. 93, 14686–14691. 10.1073/pnas.93.25.14686.

33. Jetzer, T., Studer, L., Bieri, M., Greber, U.F., and Hemmi, S. (2023). Engineered Human Adenoviruses of Species B and C Report Early, Intermediate Early, and Late Viral Gene Expression. Hum. Gene Ther. 34, 1230–1247. 10.1089/hum.2023.121.

34. Basis, A., Sharf, R., and Kleinberger, T. (2025). The adenoviral E4orf4 protein: A multifunctional protein serving as a guide for treating cancer, a multifactorial disease. Tumour Virus Res. 19, 200303. 10.1016/j.tvr.2024.200303.

35. Miron, M.-J., Blanchette, P., Groitl, P., Dallaire, F., Teodoro, J.G., Li, S., Dobner, T., and Branton, P.E. (2009). Localization and importance of the adenovirus E4orf4 protein during lytic infection. J. Virol. 83, 1689–1699. 10.1128/JVI.01703-08.

36. Marcellus, R.C., Chan, H., Paquette, D., Thirlwell, S., Boivin, D., and Branton, P.E. (2000). Induction of p53-independent apoptosis by the adenovirus E4orf4 protein requires binding to the Balpha subunit of protein phosphatase 2A. J. Virol. 74, 7869–7877. 10.1128/jvi.74.17.7869-7877.2000.

37. Cohen, P., Klumpp, S., and Schelling, D.L. (1989). An improved procedure for identifying and quantitating protein phosphatases in mammalian tissues. FEBS Lett. 250, 596–600. 10.1016/0014-5793(89)80803-8.

38. Kosulin, K., Geiger, E., Vecsei, A., Huber, W.D., Rauch, M., Brenner, E., Wrba, F., Hammer, K., Innerhofer, A., Potschger, U., et al. (2016). Persistence and reactivation of human adenoviruses in the gastrointestinal tract. Clin Microbiol Infect 22, 381 e1–8. 10.1016/j.cmi.2015.12.013.

39. Prasad, V., Suomalainen, M., Jasiqi, Y., Hemmi, S., Hearing, P., Hosie, L., Burgert, H.-G., and Greber, U.F. (2020). The UPR sensor IRE1α and the adenovirus E3-19K glycoprotein sustain persistent and lytic infections. Nat. Commun. 11, 1997. 10.1038/s41467-020-15844-2.

40. Zheng, Y., Stamminger, T., and Hearing, P. (2016). E2F/Rb Family Proteins Mediate Interferon Induced Repression of Adenovirus Immediate Early Transcription to Promote Persistent Viral Infection. PLoS Pathog 12, e1005415. 10.1371/journal.ppat.1005415.

41. Labeur, M., Refojo, D., Wölfel, B., Stalla, J., Vargas, V., Theodoropoulou, M., Buchfelder, M., Paez-Pereda, M., Arzt, E., and Stalla, G.K. (2008). Interferon-gamma inhibits cellular proliferation and ACTH production in corticotroph tumor cells through a novel janus kinases-signal transducer and activator of transcription 1/nuclear factor-kappa B inhibitory signaling pathway. J. Endocrinol. 199, 177–189. 10.1677/JOE-08-0011.

42. Walters, R.W., Grunst, T., Bergelson, J.M., Finberg, R.W., Welsh, M.J., and Zabner, J. (1999). Basolateral localization of fiber receptors limits adenovirus infection from the apical surface of airway epithelia. J. Biol. Chem. 274, 10219–10226. 10.1074/jbc.274.15.10219.

43. Sakaue-Sawano, A., Kurokawa, H., Morimura, T., Hanyu, A., Hama, H., Osawa, H., Kashiwagi, S., Fukami, K., Miyata, T., Miyoshi, H., et al. (2008). Visualizing spatiotemporal dynamics of multicellular cell-cycle progression. Cell 132, 487–498. 10.1016/j.cell.2007.12.033.

44. Robert, A., Smadja-Lamère, N., Landry, M.-C., Champagne, C., Petrie, R., Lamarche-Vane, N., Hosoya, H., and Lavoie, J.N. (2006). Adenovirus E4orf4 Hijacks Rho GTPase-dependent Actin Dynamics to Kill Cells: A Role for Endosome-associated Actin Assembly. Mol. Biol. Cell 17, 3329–3344. 10.1091/mbc.e05-12-1146.

45. Dziengelewski, C., Rodrigue, M.-A., Caillier, A., Jacquet, K., Boulanger, M.-C., Bergeman, J., Fuchs, M., Lambert, H., Laprise, P., and Richard, D.E. (2020). Adenoviral protein E4orf4 interacts with the polarity protein Par3 to induce nuclear rupture and tumor cell death. J. Cell Biol. 219.

46. Mui, M.Z., Zhou, Y., Blanchette, P., Chughtai, N., Knight, J.F., Gruosso, T., Papadakis, A.I., Huang, S., Park, M., Gingras, A.-C., et al. (2015). The Human Adenovirus Type 5 E4orf4 Protein Targets Two Phosphatase Regulators of the Hippo Signaling Pathway. J. Virol. 89, 8855–8870. 10.1128/JVI.03710-14.

47. Livne, A., Shtrichman, R., and Kleinberger, T. (2001). Caspase activation by adenovirus e4orf4 protein is cell line specific and Is mediated by the death receptor pathway. J. Virol. 75, 789–798. 10.1128/JVI.75.2.789-798.2001.

48. Schrock, M.S., Stromberg, B.R., Scarberry, L., and Summers, M.K. (2020). APC/C ubiquitin ligase: Functions and mechanisms in tumorigenesis. Semin. Cancer Biol. 67, 80–91. 10.1016/j.semcancer.2020.03.001.

49. Pesin, J.A., and Orr-Weaver, T.L. (2008). Regulation of APC/C Activators in Mitosis and Meiosis. Annu. Rev. Cell Dev. Biol. 24, 475–499. 10.1146/annurev.cellbio.041408.115949.

50. Mui, M.Z., Roopchand, D.E., Gentry, M.S., Hallberg, R.L., Vogel, J., and Branton, P.E. (2010). Adenovirus Protein E4orf4 Induces Premature APCCdc20 Activation in Saccharomyces cerevisiae by a Protein Phosphatase 2A-Dependent Mechanism. J. Virol. 84, 4798–4809. 10.1128/JVI.02434-09.

51. Tian, W., Li, B., Warrington, R., Tomchick, D.R., Yu, H., and Luo, X. (2012). Structural analysis of human Cdc20 supports multisite degron recognition by APC/C. Proc. Natl. Acad. Sci. U. S. A. 109, 18419–18424. 10.1073/pnas.1213438109.

52. Pesin, J.A., and Orr-Weaver, T.L. (2008). Regulation of APC/C activators in mitosis and meiosis. Annu. Rev. Cell Dev. Biol. 24, 475–499. 10.1146/annurev.cellbio.041408.115949.

53. Hyun, S.-Y., Sarantuya, B., Lee, H.-J., and Jang, Y.-J. (2013). APC/C(Cdh1)-dependent degradation of Cdc20 requires a phosphorylation on CRY-box by Polo-like kinase-1 during somatic cell cycle. Biochem. Biophys. Res. Commun. 436, 12–18. 10.1016/j.bbrc.2013.04.073.

54. Sakaue-Sawano, A., Yo, M., Komatsu, N., Hiratsuka, T., Kogure, T., Hoshida, T., Goshima, N., Matsuda, M., Miyoshi, H., and Miyawaki, A. (2017). Genetically Encoded Tools for Optical Dissection of the Mammalian Cell Cycle. Mol. Cell 68, 626–640.e5. 10.1016/j.molcel.2017.10.001.

55. Ge, S., Skaar, J.R., and Pagano, M. (2009). APC/C- and Mad2-mediated degradation of Cdc20 during spindle checkpoint activation. Cell Cycle Georget. Tex 8, 167–171. 10.4161/cc.8.1.7606.

56. Garcia-Gomez, A., Quwaider, D., Canavese, M., Ocio, E.M., Tian, Z., Blanco, J.F., Berger, A.J., Ortiz-de-Solorzano, C., Hernández-Iglesias, T., Martens, A.C.M., et al. (2014). Preclinical activity of the oral proteasome inhibitor MLN9708 in Myeloma bone disease. Clin. Cancer Res. Off. J. Am. Assoc. Cancer Res. 20, 1542–1554. 10.1158/1078-0432.CCR-13-1657.

57. Mackay, D.R., Makise, M., and Ullman, K.S. (2010). Defects in nuclear pore assembly lead to activation of an Aurora B-mediated abscission checkpoint. J. Cell Biol. 191, 923–931. 10.1083/jcb.201007124.

58. Nguyen, H.G., Chinnappan, D., Urano, T., and Ravid, K. (2005). Mechanism of Aurora-B Degradation and Its Dependency on Intact KEN and A-Boxes: Identification of an Aneuploidy-Promoting Property. Mol. Cell. Biol. 25, 4977–4992. 10.1128/MCB.25.12.4977-4992.2005.

59. Graham, F.L., Smiley, J., Russell, W.C., and Nairn, R. (1977). Characteristics of a human cell line transformed by DNA from human adenovirus type 5. J. Gen. Virol. 36, 59–74. 10.1099/0022-1317-36-1-59.

60. Sackton, K.L., Dimova, N., Zeng, X., Tian, W., Zhang, M., Sackton, T.B., Meaders, J., Pfaff, K.L., Sigoillot, F., Yu, H., et al. (2014). Synergistic blockade of mitotic exit by two chemical inhibitors of the APC/C. Nature 514, 646–649. 10.1038/nature13660.

61. Barr, F.A., and Gruneberg, U. (2007). Cytokinesis: placing and making the final cut. Cell 131, 847–860. 10.1016/j.cell.2007.11.011.

62. Ettinger, A.W., Wilsch-Brauninger, M., Marzesco, A.M., Bickle, M., Lohmann, A., Maliga, Z., Karbanova, J., Corbeil, D., Hyman, A.A., and Huttner, W.B. (2011). Proliferating versus differentiating stem and cancer cells exhibit distinct midbody-release behaviour. Nat Commun 2, 503. 10.1038/ncomms1511.

63. Sakuma, T., Nakade, S., Sakane, Y., Suzuki, K.-I.T., and Yamamoto, T. (2016). MMEJ-assisted gene knock-in using TALENs and CRISPR-Cas9 with the PITCh systems. Nat. Protoc. 11, 118–133. 10.1038/nprot.2015.140.

64. Adams, R.R., Maiato, H., Earnshaw, W.C., and Carmena, M. (2001). Essential roles of Drosophila inner centromere protein (INCENP) and aurora B in histone H3 phosphorylation, metaphase chromosome alignment, kinetochore disjunction, and chromosome segregation. J. Cell Biol. 153, 865–880. 10.1083/jcb.153.4.865.

65. Cooke, C.A., Heck, M.M., and Earnshaw, W.C. (1987). The inner centromere protein (INCENP) antigens: movement from inner centromere to midbody during mitosis. J. Cell Biol. 105, 2053–2067. 10.1083/jcb.105.5.2053.

66. Carmena, M., Wheelock, M., Funabiki, H., and Earnshaw, W.C. (2012). The chromosomal passenger complex (CPC): from easy rider to the godfather of mitosis. Nat. Rev. Mol. Cell Biol. 13, 789–803. 10.1038/nrm3474.

67. Shitova, M., Alpeeva, E., and Vorotelyak, E. (2024). Review of hTERT-Immortalized Cells: How to Assess Immortality and Confirm Identity. Int. J. Mol. Sci. 25, 13054. 10.3390/ijms252313054.

68. Hohlweg, U., Dorn, A., Hösel, M., Webb, D., Buettner, R., and Doerfler, W. (2004). Tumorigenesis by adenovirus type 12 in newborn Syrian hamsters. Curr. Top. Microbiol. Immunol. 273, 215–244. 10.1007/978-3-662-05599-1_7.

69. Kosulin, K., Haberler, C., Hainfellner, J.A., Amann, G., Lang, S., and Lion, T. (2007). Investigation of adenovirus occurrence in pediatric tumor entities. J. Virol. 81, 7629–7635. 10.1128/JVI.00355-07.

70. Shtrichman, R., Sharf, R., Barr, H., Dobner, T., and Kleinberger, T. (1999). Induction of apoptosis by adenovirus E4orf4 protein is specific to transformed cells and requires an interaction with protein phosphatase 2A. Proc. Natl. Acad. Sci. U. S. A. 96, 10080–10085. 10.1073/pnas.96.18.10080.

71. Thaker, S.K., Ch’ng, J., and Christofk, H.R. (2019). Viral hijacking of cellular metabolism. BMC Biol. 17, 59. 10.1186/s12915-019-0678-9.

72. Pines, J. (2011). Cubism and the cell cycle: the many faces of the APC/C. Nat. Rev. Mol. Cell Biol. 12, 427–438. 10.1038/nrm3132.

73. Labit, H., Fujimitsu, K., Bayin, N.S., Takaki, T., Gannon, J., and Yamano, H. (2012). Dephosphorylation of Cdc20 is required for its C-box-dependent activation of the APC/C. EMBO J. 31, 3351–3362. 10.1038/emboj.2012.168.

74. Yamano, H. (2019). APC/C: current understanding and future perspectives. Preprint at F1000Research, 10.12688/f1000research.18582.1 https://doi.org/10.12688/f1000research.18582.1.

75. Hein, J.B., Hertz, E.P.T., Garvanska, D.H., Kruse, T., and Nilsson, J. (2017). Distinct kinetics of serine and threonine dephosphorylation are essential for mitosis. Nat. Cell Biol. 19, 1433–1440. 10.1038/ncb3634.

76. Cundell, M.J., Hutter, L.H., Nunes Bastos, R., Poser, E., Holder, J., Mohammed, S., Novak, B., and Barr, F.A. (2016). A PP2A-B55 recognition signal controls substrate dephosphorylation kinetics during mitotic exit. J. Cell Biol. 214, 539–554. 10.1083/jcb.201606033.

77. Fang, G., Yu, H., and Kirschner, M.W. (1998). Direct binding of CDC20 protein family members activates the anaphase-promoting complex in mitosis and G1. Mol. Cell 2, 163–171. 10.1016/s1097-2765(00)80126-4.

78. Chen, O.J., Feng, C.Y., Ladak, R.J., Crosby, M.A., Russo, M. de S.T., Wang, Y., Blanchette, P., Liang, Y., Sharon, D.M., Moustafa-Kamal, M., et al. (2026). Viral inhibition of the anaphase promoting complex enhances replication by elevating nucleotide pools. Preprint at bioRxiv, 10.64898/2026.01.26.701660 https://doi.org/10.64898/2026.01.26.701660.

79. Wolf, A., Keil, R., Götzl, O., Mun, A., Schwarze, K., Lederer, M., Hüttelmaier, S., and Hatzfeld, M. (2006). The armadillo protein p0071 regulates Rho signalling during cytokinesis. Nat. Cell Biol. 8, 1432–1440. 10.1038/ncb1504.

80. Prasad, V., Cerikan, B., Stahl, Y., Kopp, K., Magg, V., Acosta-Rivero, N., Kim, H., Klein, K., Funaya, C., Haselmann, U., et al. (2023). Enhanced SARS-CoV-2 entry via UPR-dependent AMPK-related kinase NUAK2. Mol. Cell 83, 2559–2577. 10.1016/j.molcel.2023.06.020.

81. Suomalainen, M., Prasad, V., Kannan, A., and Greber, U.F. (2020). Cell-to-cell and genome-to-genome variability of adenovirus transcription tuned by the cell cycle. J Cell Sci 134. 10.1242/jcs.252544.

82. Stringer, C., Wang, T., Michaelos, M., and Pachitariu, M. (2021). Cellpose: a generalist algorithm for cellular segmentation. Nat Methods 18, 100–106. 10.1038/s41592-020-01018-x.

83. Carpenter, A.E., Jones, T.R., Lamprecht, M.R., Clarke, C., Kang, I.H., Friman, O., Guertin, D.A., Chang, J.H., Lindquist, R.A., Moffat, J., et al. (2006). CellProfiler: image analysis software for identifying and quantifying cell phenotypes. Genome Biol 7, R100. 10.1186/gb-2006-7-10-r100.

84. Dietz, C., and Berthold, M.R. (2016). KNIME for Open-Source Bioimage Analysis: A Tutorial. Adv Anat Embryol Cell Biol 219, 179–197. 10.1007/978-3-319-28549-8_7.

85. Müller, T., Kalxdorf, M., Longuespée, R., Kazdal, D.N., Stenzinger, A., and Krijgsveld, J. (2020). Automated sample preparation with SP3 for low-input clinical proteomics. Mol. Syst. Biol. 16, e9111. 10.15252/msb.20199111.

86. Tyanova, S., Temu, T., and Cox, J. (2016). The MaxQuant computational platform for mass spectrometry-based shotgun proteomics. Nat. Protoc. 11, 2301–2319. 10.1038/nprot.2016.136.

87. Schwanhäusser, B., Busse, D., Li, N., Dittmar, G., Schuchhardt, J., Wolf, J., Chen, W., and Selbach, M. (2011). Global quantification of mammalian gene expression control. Nature 473, 337–342. 10.1038/nature10098.

