## Supplementary figures for "Viral rewiring of APC/C-CDC20 drives Aurora B hyperubiquitination, mitotic regression and polyploidy"

**This PDF file includes:**

Figures S1 to S11
Legends for Movie S1-S7
Legends for Table S1
Table of reagents
References

**Other Supplementary Materials for this manuscript include the following:**

| | |
|---|---|
| Table S1 | Proteome analysis |
| Movies S1-S7 | Movies S1-S7 |

#### Supplementary Figures

### Figure S1

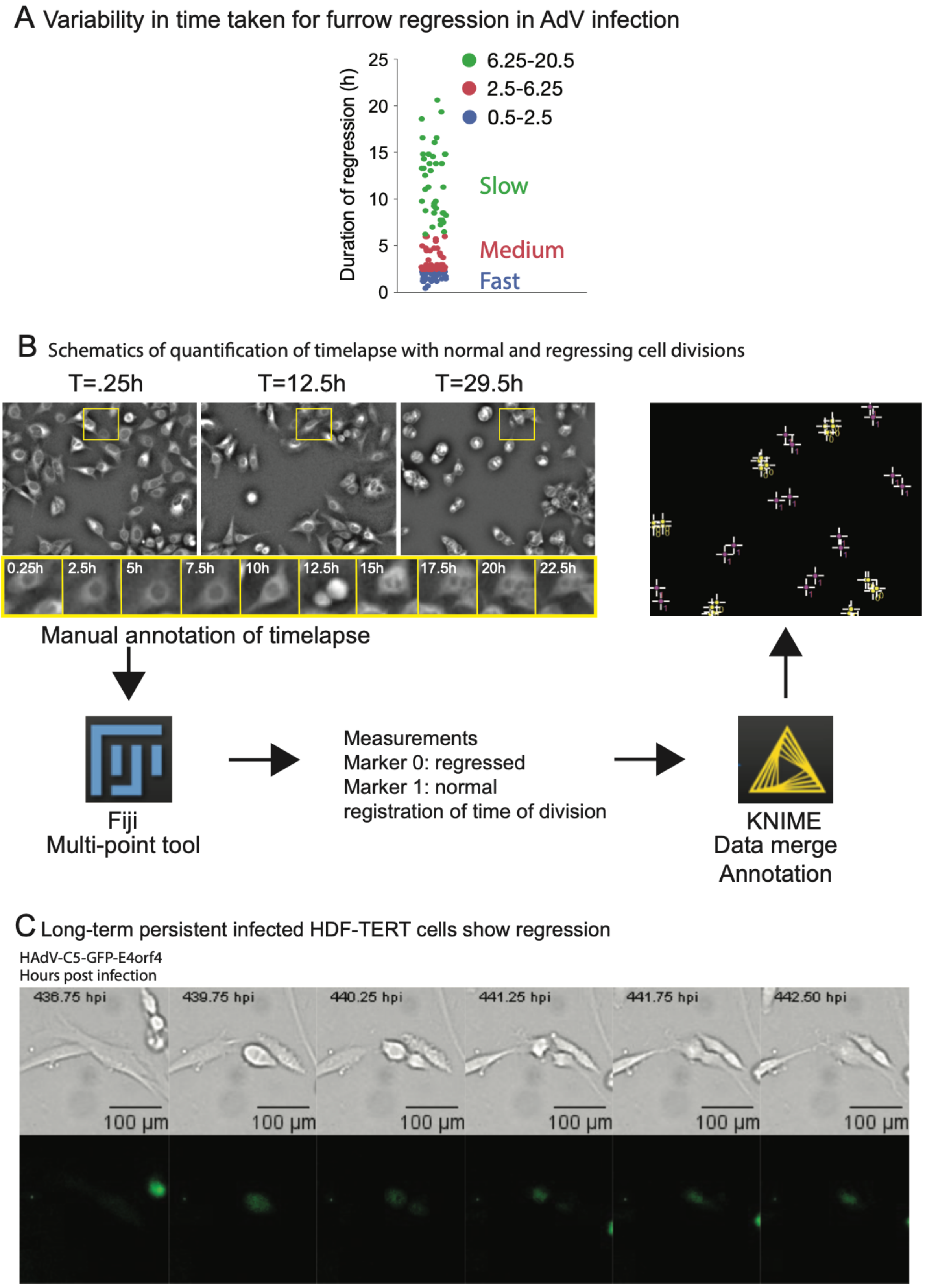

**Figure S1. Variability, quantification strategy, and persistence of cleavage furrow regression during AdV infection**

(A) Duration between cleavage furrow ingression to regression following AdV infection. Each dot represents one regression event, plotted as the time elapsed between initial furrow ingression and completion of regression. Regression events were grouped into fast (0.5–2.5 h), medium (2.5–6.25 h), and slow (6.25–20.5h) categories, illustrating substantial heterogeneity in regression kinetics.

(B) Workflow for the quantification of normal and regressing cell divisions from long-term time-lapse imaging. Representative A549-Sec61ß-GFP fluorescent images show tracking of individual cells across imaging time points, with zoomed in insets highlighting division progression. Cell divisions were manually annotated using the Fiji “multipoint tool” to assign division outcome (regressed or normal) and division timing, followed by data merging and annotation using KNIME data analytics for downstream analysis.

(C) Representative long-term time-lapse images showing cleavage furrow regression in persistently infected HDF-TERT cells expressing GFP-E4orf4. Phase-contrast (top) and GFP fluorescence (bottom) images demonstrate that regression events in persistent AdV infection model can occur several hundred hours after infection, indicating sustained cytokinesis defects during persistent adenovirus infection. Scale bars, 100 µm.

Figure S2

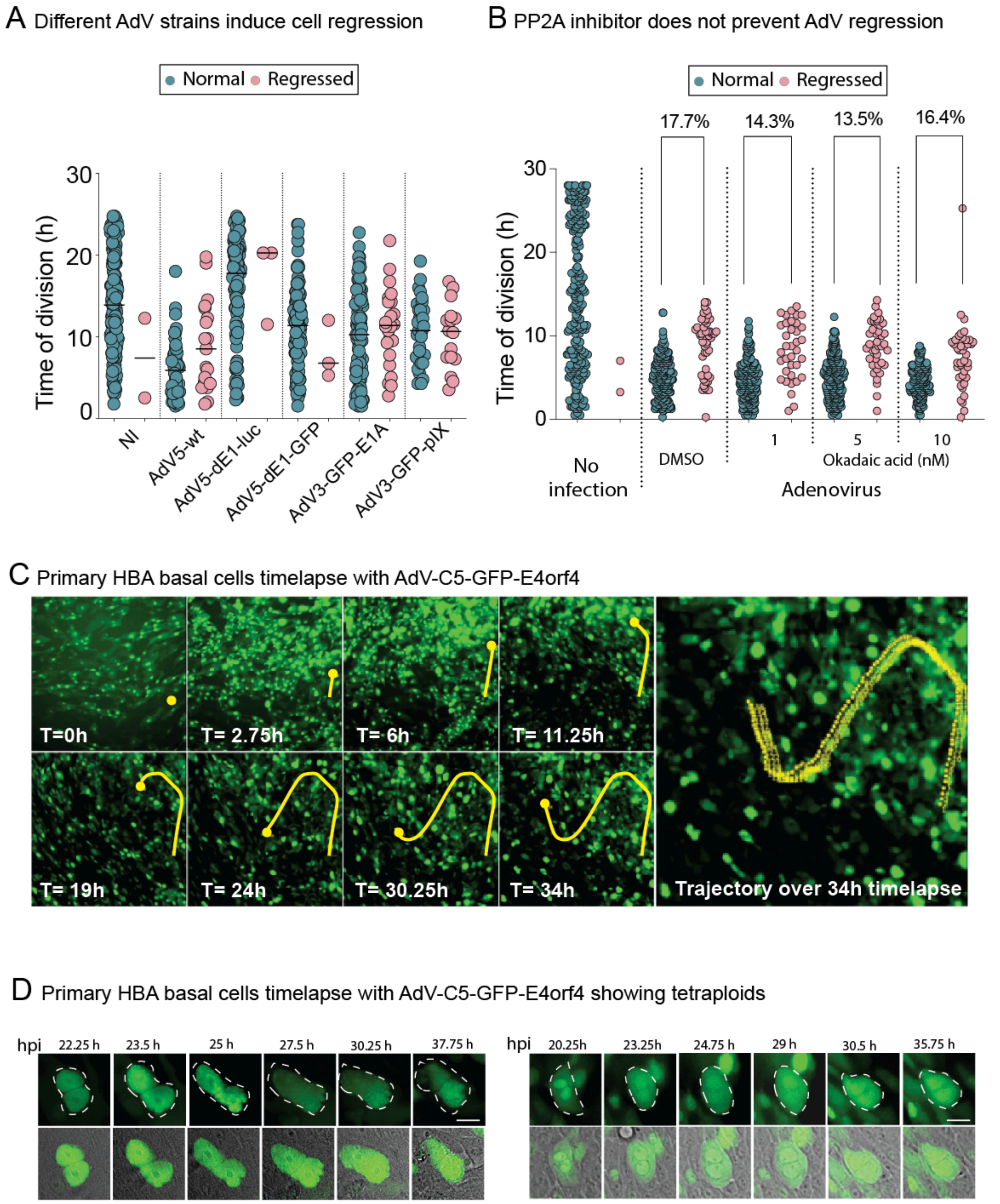


**Figure S2. Cleavage furrow regression occurs across adenovirus strains and is not blocked by PP2A inhibition**

(A) Cell division timing and cytokinesis outcome (normal or regression) following infection with different AdV strains. Scatter plots show the time of division for non-infected (NI) cells and cells infected with AdV5-wt, AdV5-dE1-luc, AdV5-dE1-GFP, AdV3-GFP-E1A, and AdV3-GFP-pIX. Each dot represents one cell division, classified as normal (blue) or regressed (pink), indicating that cleavage furrow regression requires AdV gene expression, and is observed across multiple adenovirus serotypes and genetic backgrounds.

(B) Effect of PP2A inhibition on AdV-induced cleavage furrow regression. Cells were infected with adenovirus and treated with DMSO or increasing concentrations of okadaic acid (1, 5, or 10 nM), and division outcomes were quantified by live-cell imaging. Scatter plots show single-cell division times for normal (teal) and regressed (pink) divisions, with the percentage of regressed divisions indicated above each condition, demonstrating that PP2A inhibition does not prevent AdV-induced regression.

(C) High migratory nature of primary HBA basal cells infected with AdV-C5–GFP-E4orf4 during long-term time-lapse imaging. A representative infected cell (yellow dot) is shown. Due to the high migratory nature of primary HBA basal cells, sustained single-cell tracking across the full imaging window was feasible only for a limited subset of cells.

(D) Binucleated primary HBA basal cells infected with AdV-C5–GFP-E4orf4. Two representative cells infected with AdV-C5-GFP-E4orf4 are shown with binucleate condition. Scale bar, 10μm.

Figure S3

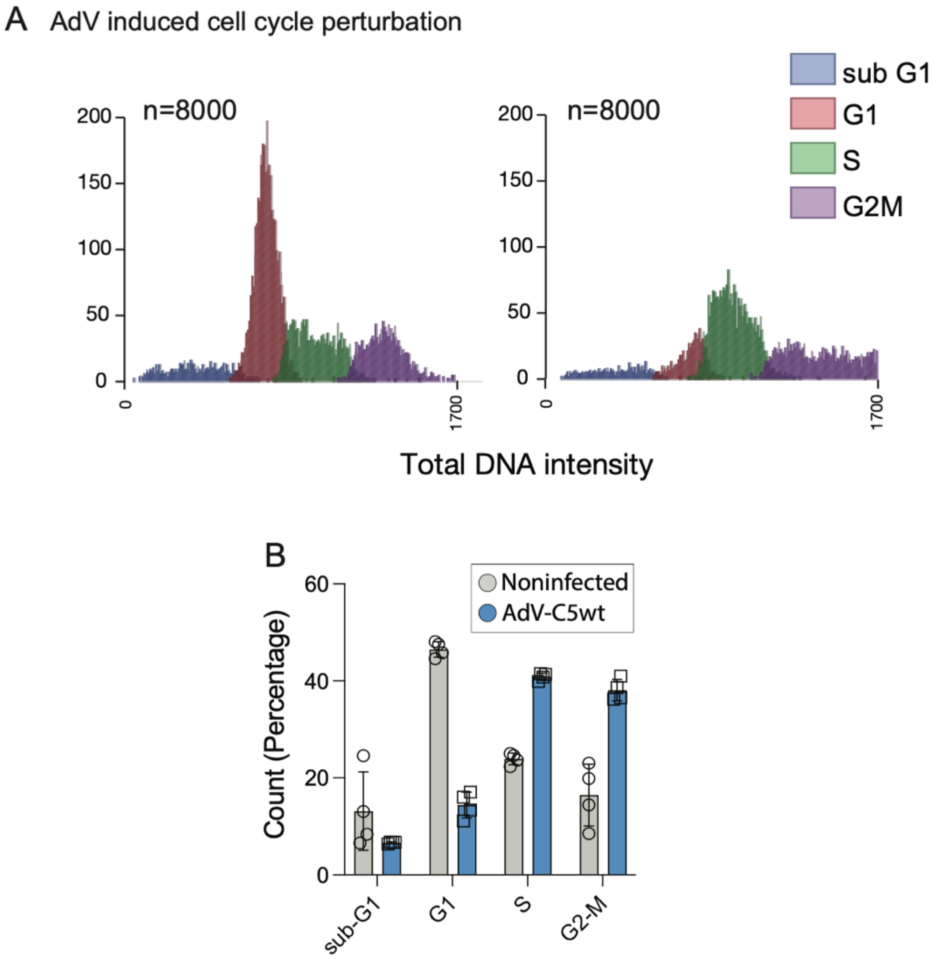


**Figure S3. Adenovirus infection perturbs cell cycle distribution of proliferating cells**

(A) Cell cycle distribution of non-infected and AdV-C5-infected cells assessed by total DNA content analysis. Representative DNA intensity histograms (left) show classification of cells into sub-G1, G1, S, and G2/M populations (n = 8,000 cells per condition).

(B) Quantification (right) shows the percentage of cells in each cell cycle phase for non-infected and AdV-C5-infected populations, demonstrating AdV-induced increase in cells in S and G2/M cell cycle stages.

Figure S4

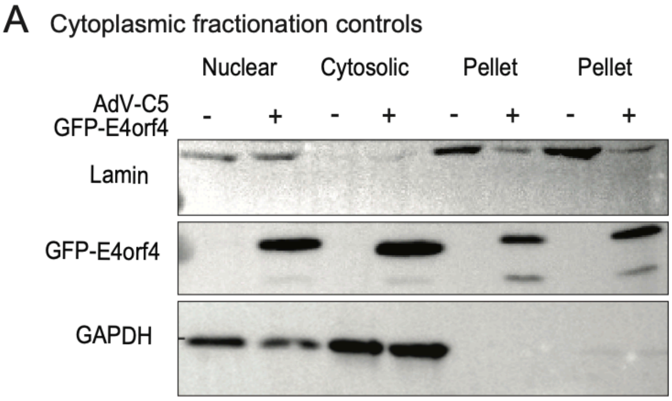


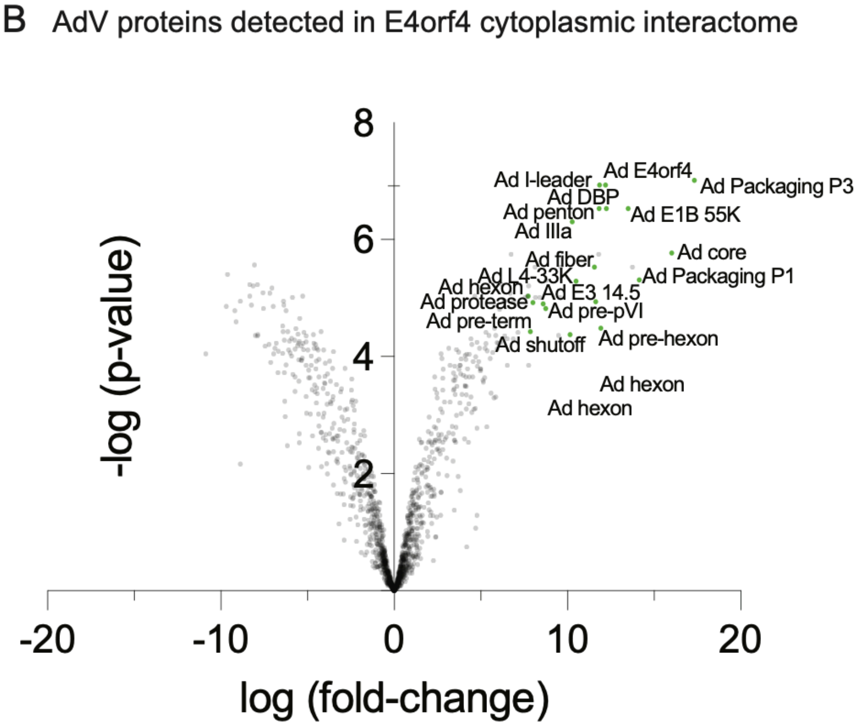


**Figure S4. Validation of cytoplasmic fractionation and identification of adenoviral proteins in the E4orf4 cytoplasmic interactome**

(A) Validation of cytoplasmic fractionation used for E4orf4 interactome analysis. Immunoblot analysis of nuclear, cytosolic, and pellet fractions from cells infected with AdV-C5 expressing

GFP-E4orf4 or control cells. Lamin and GAPDH were used as markers for nuclear and cytosolic compartments, respectively, confirming efficient fractionation and minimal cross-contamination. GFP-E4orf4 is detected in nuclear, cytosolic, and pellet fractions in infected cells.

(B) Identification of AdV proteins associated with the E4orf4 cytoplasmic interactome. Volcano plot showing AdV proteins significantly enriched in GFP-E4orf4 immunoprecipitates from the cytoplasmic fraction, plotted as log fold-change versus –log(p-value). Several early and late viral proteins, including structural and packaging components, are detected, indicating broad association of E4orf4 with viral proteins during infection.

Figure S5

A Recombinant protein expression and purification

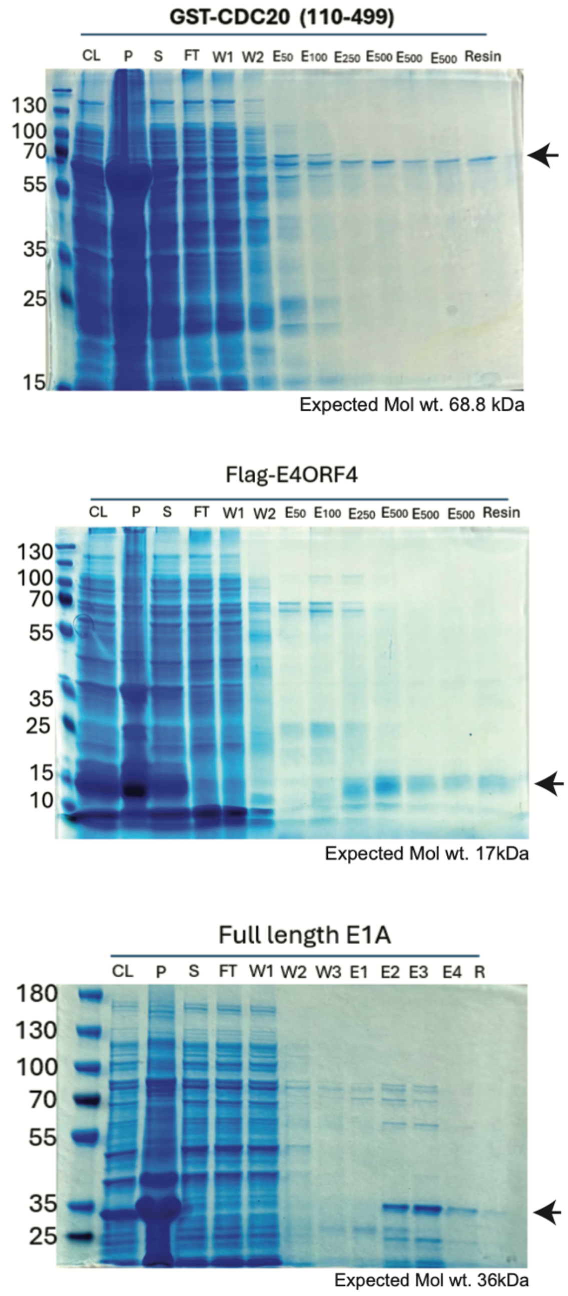


**Figure S5. Recombinant protein purification for CDC20, E4orf4, and E1A.**

Coomassie-stained SDS–PAGE analysis of recombinant proteins used in *in vitro* binding assays. Representative purification profiles are shown for GST-CDC20 (amino acids 110–499), FLAG–E4orf4, and full-length E1A. Lanes indicate cleared lysate (CL), pellet (P), supernatant (S), flow-through (FT), wash fractions (W), elution fractions (E), and resin, demonstrating enrichment of the indicated recombinant proteins to near homogeneity.

Figure S6

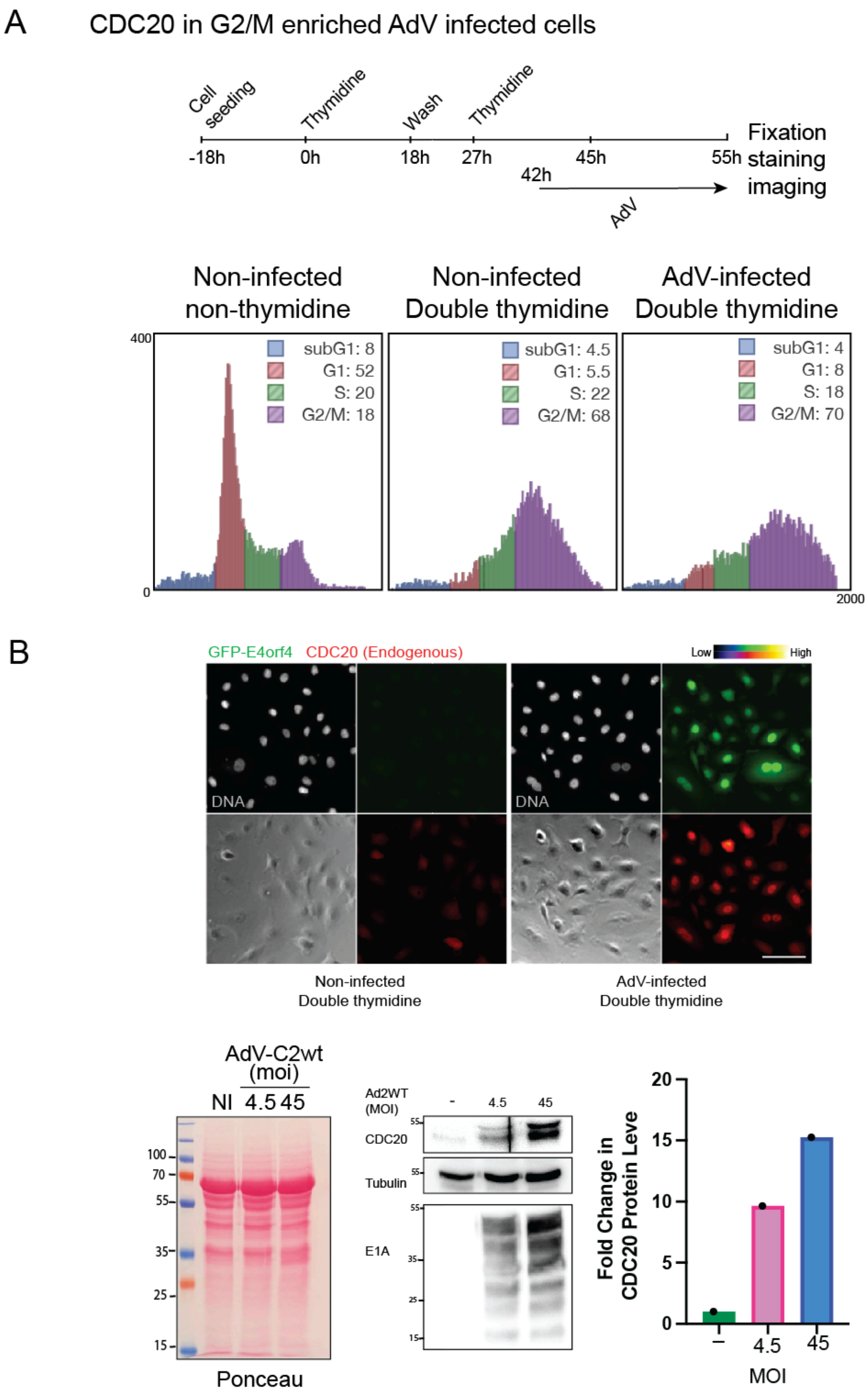


**Figure S6. CDC20 accumulation in G2/M-enriched AdV-infected cells**

(A) Experimental scheme and cell cycle enrichment strategy used to assess CDC20 levels in AdV-infected cells. Cells were synchronized using a double-thymidine block, infected with adenovirus, and collected for fixation, staining, and imaging at the indicated time points.

Representative DNA content histograms show cell cycle distributions for non-infected asynchronous cells, non-infected double-thymidine-treated cells, and AdV-infected double-thymidine-treated cells, demonstrating effective enrichment of the G2/M population following synchronization and infection.

(B) Immunofluorescence analysis of endogenous CDC20 and GFP-E4orf4 expression in synchronized cells. Representative images show DNA staining, GFP-E4orf4 fluorescence, and endogenous CDC20 signal in non-infected and AdV-infected cells following double-thymidine synchronization. Heat map representation indicates relative CDC20 intensity levels, revealing elevated CDC20 signal in AdV-infected cells compared to non-infected controls. Scale bar, 100μm.

(C) Immunoblot analysis of CDC20 protein levels following adenovirus infection at increasing multiplicities of infection (MOI). Ponceau staining is shown as a loading control, and immunoblots for CDC20, tubulin, and the viral protein E1A confirm infection efficiency. Quantification (right) shows fold change in CDC20 protein levels relative to non-infected cells, indicating a dose-dependent increase in CDC20 abundance upon AdV infection.

Figure S7

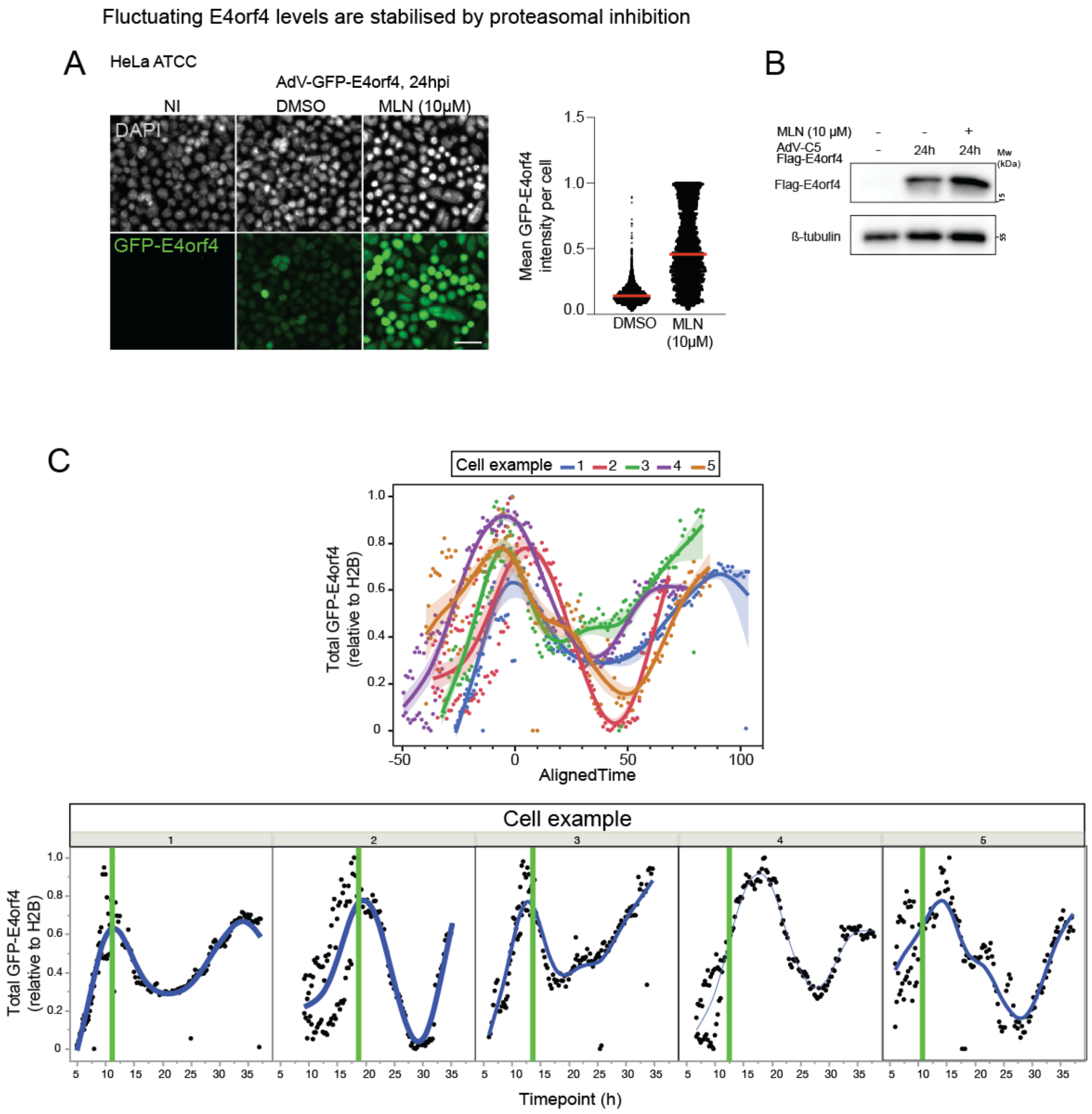


**Figure S7. Proteasomal inhibition stabilizes fluctuating E4orf4 protein levels during adenovirus infection**

(A) Immunofluorescence analysis of GFP-E4orf4 expression in HeLa cells under proteasome-inhibited conditions. Non-infected (NI) cells and cells infected with AdV-GFP-E4orf4 for 24 h were treated with DMSO or the proteasome inhibitor MLN (10 μM) as indicated. Representative images show DAPI-stained nuclei and GFP-E4orf4 fluorescence, with single-cell quantification of mean GFP-E4orf4 intensity per cell demonstrating increased and stabilized E4orf4 levels upon proteasome inhibition. Scale bar, 50μm.

(B) Immunoblot analysis confirming proteasome-dependent regulation of E4orf4 protein levels. Cells infected with AdV-C5 expressing FLAG-E4orf4 were treated with DMSO or MLN (10 μM) for 24 h, and whole-cell lysates were analyzed by immunoblotting for FLAG-E4orf4. β-tubulin serves as a loading control, showing increased accumulation of E4orf4 upon proteasome inhibition.

(C) Single-cell time-lapse analysis of GFP-E4orf4 dynamics during adenovirus infection. Top, normalized GFP-E4orf4 intensity traces from five representative cells aligned in time, illustrating oscillatory E4orf4 expression dynamics. Bottom, individual GFP-E4orf4 intensity traces plotted over time for the same cells, with vertical green lines indicating the time of cleavage furrow regression, demonstrating that regression events occur proximal to peaks in E4orf4 abundance.

### Figure S8

A E4orf4 expressed from native E4 promoter fluctuate in infection

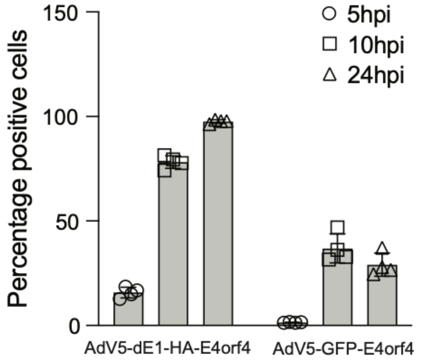


B GFP-E4orf4 expression from E4 promoter in E1A expressing cell lines

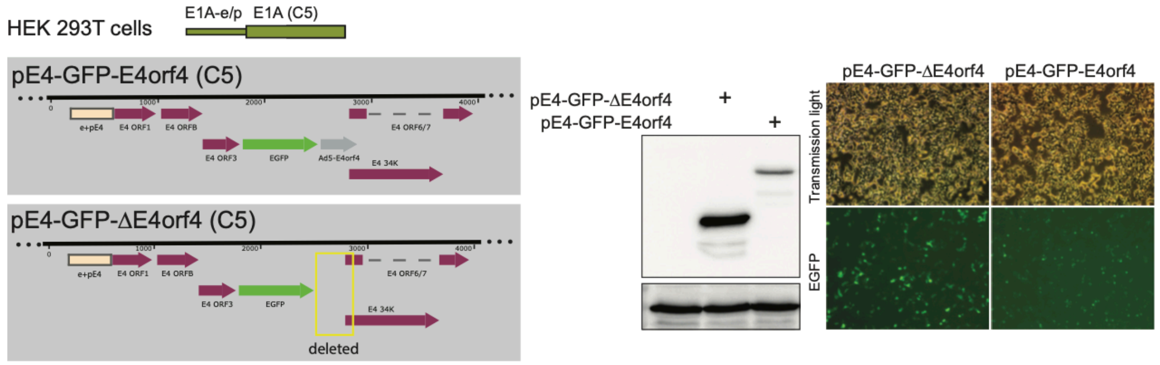


C Apcin and proTAME effect on E4orf4-CDC20-AurB complex

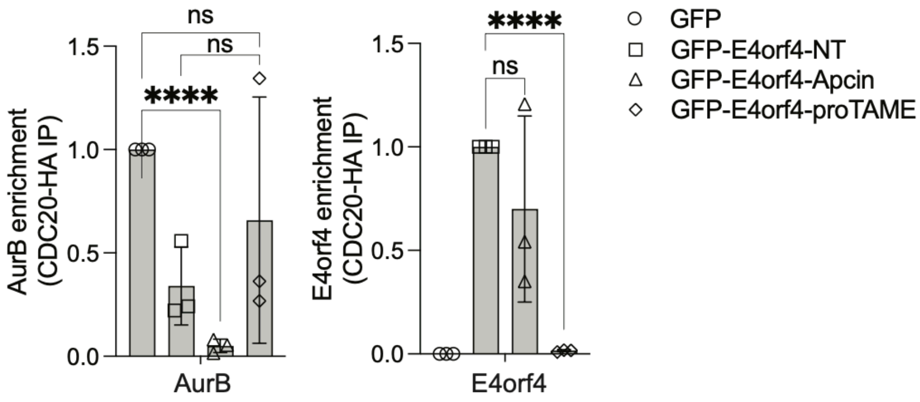


**Figure S8. Fluctuating expression of E4orf4 from the native E4 promoter during AdV infection**

(A) Quantification of E4orf4-positive cells during AdV infection when E4orf4 is expressed from its native E4 promoter. The percentage of E4orf4-positive cells was measured at 5, 10, and 24 hours post infection (hpi) for AdV5-dE1-HA-E4orf4 and AdV5-GFP-E4orf4. Data show timedependent fluctuations in the fraction of E4orf4-expressing cells, indicating dynamic regulation of E4orf4 expression during infection.

(B) E4 promoter–driven expression of GFP–E4orf4 in E1A-expressing cell lines. Schematic representation of pE4-GFP-E4orf4 (C5) and pE4-GFP-ΔE4orf4 (C5) constructs used to assess E4 promoter activity in HEK293T cells, which constitutively express E1A. Immunoblot analysis confirms GFP–E4orf4 expression from the pE4 construct, while deletion of E4orf4 abrogates expression. Representative transmission light and EGFP fluorescence images show GFP signal in cells transfected with pE4-GFP-E4orf4 but not with the ΔE4orf4 control, demonstrating E1A-dependent activation of the E4 promoter.

(C) Quantification of AurB and E4orf4 enrichment with CDC20-HA immunoprecipitation in the presence of Apcin and proTAME inhibitors. Related to Figure 5G. Significance was calculated using ordinary one-way ANOVA. **** = $p<0.0001$.

### Figure S9

A mScarlet knockin in AuroraB using MMEJ

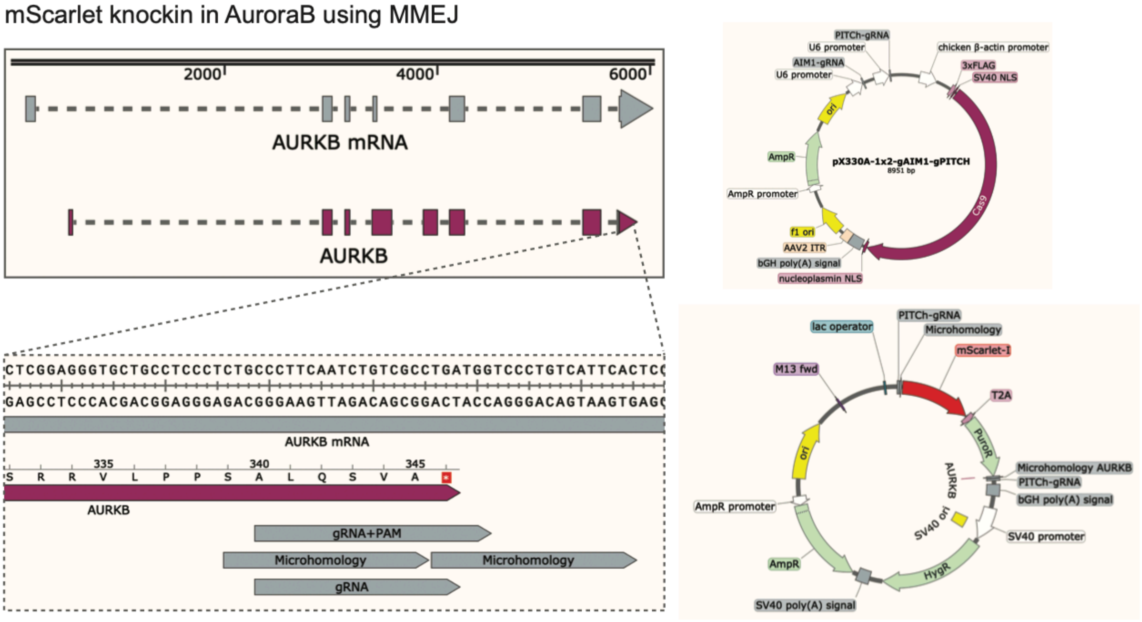


B Representative images of AurB-mScar localisation in HEK293T knock-in cells

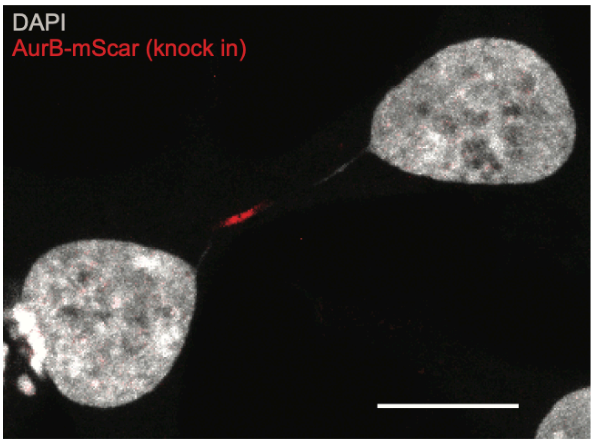


**Figure S9. Generation and validation of mScarlet-Aurora B knock-in cells**

(A) Strategy for endogenous tagging of Aurora B with mScarlet using microhomology-mediated end joining (MMEJ). Schematic illustrates the AURKB genomic locus, the CRISPR/Cas9 target site, and the donor construct containing mScarlet flanked by microhomology arms. Plasmid maps of the gRNA/Cas9 and mScarlet donor vectors used for genome editing are shown, highlighting the design used to generate C-terminally tagged Aurora B.

(B) Representative image showing mScarlet–Aurora B knock-in cells showing the typical Aurora B expression pattern at the midbody, confirming correct localization and expression of the knock-in tagged protein.

### Figure S10

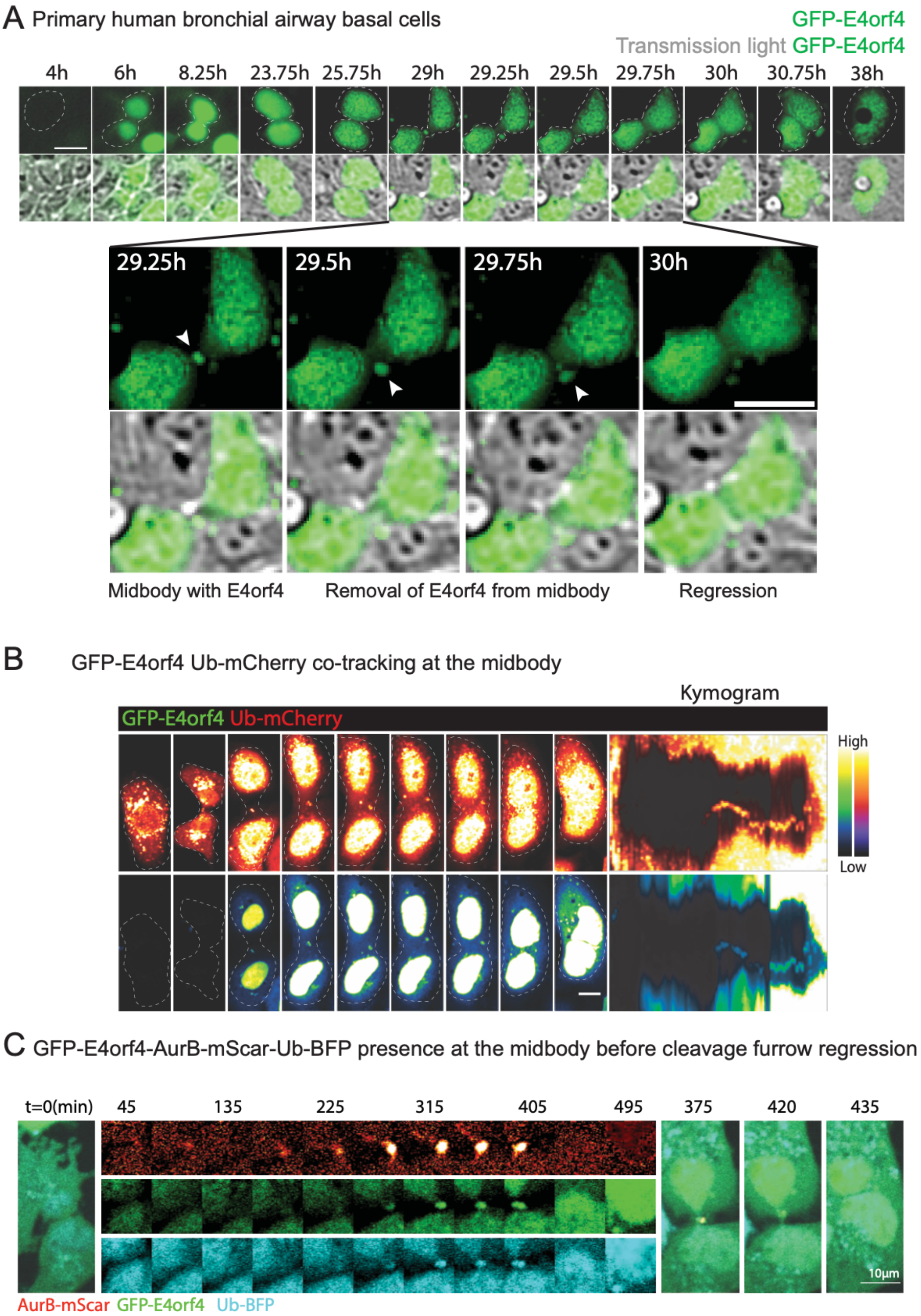
A Primary human bronchial airway basal cells
GFP-E4orf4
Transmission light GFP-E4orf4
4h
6h
8.25h
23.75h
25.75h
29h
29.25h
29.5h
29.75h
30h
30.75h
38h
29.25h
29.5h
29.75h
30h
Midbody with E4orf4
Removal of E4orf4 from midbody
Regression
B GFP-E4orf4 Ub-mCherry co-tracking at the midbody
Kymogram
GFP-E4orf4 Ub-mCherry
High
Low
C GFP-E4orf4-AurB-mScar-Ub-BFP presence at the midbody before cleavage furrow regression
t=0(min)
45
135
225
315
405
495
375
420
435
10μm
AurB-mScar GFP-E4orf4 Ub-BFP

**Figure S10. Dynamics of E4orf4 and ubiquitin at the midbody prior to cleavage furrow regression in primary and cancer cells**

(A) Time-lapse imaging of human primary bronchial airway basal cells infected with AdV-C5-GFP-E4orf4. Representative image sequences show the GFP-E4orf4 signal accumulation at the midbody region of a cell in cytokinesis, followed by regression. Inset shows zoomed out image sequences where GFP-E4orf4 puncta is seen at the midbody (highlighted with white arrows) and midbody collapse post GFP-E4orf4 dislocation from the midbody region. Scale bar, 10 µm.

(B) Time-lapse imaging of GFP-E4orf4 and Ub-mCherry co-localization at the midbody during cytokinesis in AdV-infected cells. Representative image sequences show the spatial and temporal accumulation of ubiquitin and E4orf4 at the midbody region, with corresponding kymograms illustrating signal intensity dynamics over time. Color scales indicate relative fluorescence intensity (low to high), highlighting coordinated enrichment and persistence of both signals at the midbody. Scale bar, 10µm.

(C) Simultaneous visualization of GFP-E4orf4, Aurora B-mScarlet, and Ub-BFP at the midbody prior to cleavage furrow regression. Time-lapse frames show Aurora B, E4orf4, and ubiquitin signals co-existing at the midbody before regression onset, with enlarged views illustrating signal overlap at later time points. Scale bar, 10 µm.

Figure S11

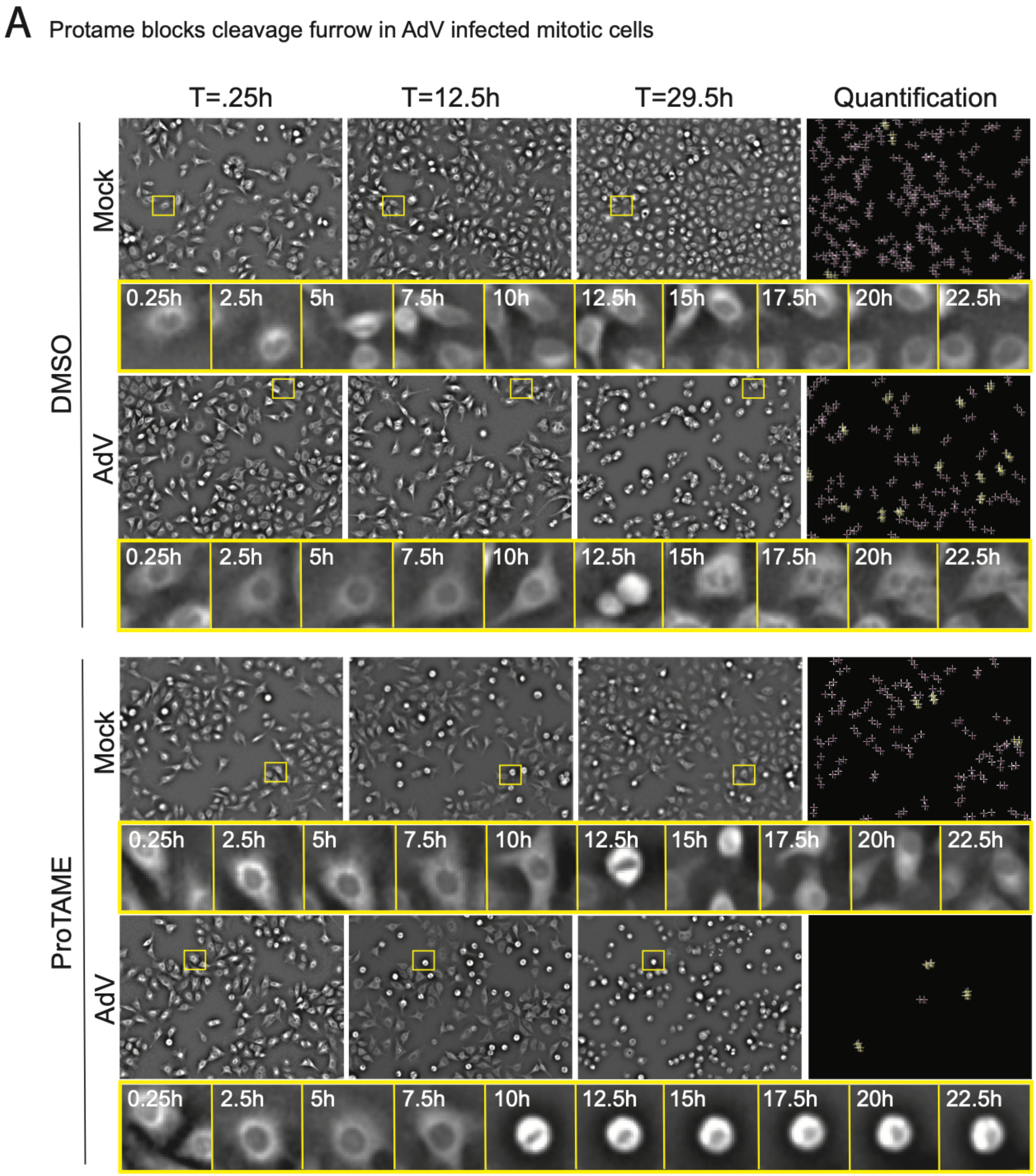


**Figure S11. ProTAME inhibits cytokinesis and cleavage furrow regression in AdV-infected mitotic cells**

(A) Time-lapse analysis of mitotic progression in mock- or AdV-infected cells treated with DMSO or ProTAME. Representative phase-contrast images show cells at indicated time points, with yellow boxes marking cells tracked over time and magnified below to illustrate division progression. In DMSO-treated AdV-infected cells, cleavage furrow ingression is followed by regression, whereas ProTAME-treated AdV-infected cells fail to undergo furrow regression and remain arrested in a rounded mitotic state. Right panels show automated quantification of division outcomes for each condition.

#### Supplementary Movies

**Movie S1: Nuclear membrane rearrangement in AdV-infected polyploid cells.**
Time-lapse imaging of HeLa cells stably expressing mRFP-Lap2β (nuclear envelope marker) following infection with AdV-C5-wt. An infected cell progresses through mitosis, undergoes cleavage furrow ingression, but fails abscission, resulting in furrow regression and formation of a tetraploid cell. Following regression, the two daughter nuclei remain initially distinct and are surrounded by Lap2β-positive nuclear envelopes. Over time, these nuclei undergo dynamic repositioning and gradual nuclear rearrangement, leading to the formation of a single, congressed nuclear structure within the tetraploid cell. Images were acquired at time intervals of 15min with timestamps are in hours:minutes. Scale bar, 10 µm.

**Movie S2-3: AurB-mScarlet knock-in cells show AurB dynamics during AdV infection.**
Time-lapse imaging of AurB-mScarlet knock-in HEK293T cells during AdV-C5-GFP-E4orf4 infection. AurB localizes normally to chromosomes and the midbody during mitotic progression (Movie 2) but displays altered spatial and temporal dynamics in infected cells (Movie 3). In cells undergoing cytokinetic failure, AurB persists at the midbody and shows abnormal redistribution coincident with cleavage furrow regression and tetraploid cell formation. Images were acquired at the indicated time intervals; timestamps are shown as minutes:seconds. Scale bar, 10µm.

**Movie S4-5: GFP-E4orf4 and Ub-mCherry co-tracking in midbody of AdV infected cells.**
Time-lapse imaging of cells infected with AdV-C5-GFP-E4orf4 infected mCherry-tagged ubiquitin expressing A549 cells. GFP-E4orf4 accumulates at the cytokinetic midbody, where it co-localizes and co-tracks with ubiquitin signals during late cytokinesis (Movie 4). Enrichment of ubiquitin at the midbody coincides temporally with E4orf4 residency and precedes cleavage furrow regression and midbody collapse, leading to tetraploid cell formation (Movie 5). Images were acquired with 15min time-resolution and timestamps are shown as hours:minutes. Scale bar, 10 µm.

**Movie S6: Increased ubiquitin residence at the midbody of AdV infected cells prior to cleavage furrow regression.**
Time-lapse imaging of cells expressing mCherry-tagged ubiquitin under non-infected and AdV-C5-wt-infected conditions. In non-infected cells, ubiquitin transiently localizes to the midbody and is rapidly cleared prior to abscission. In contrast, adenovirus-infected cells show extended retention of ubiquitin at the midbody during late cytokinesis. This increased ubiquitin residence temporally precedes cleavage furrow regression and midbody collapse, culminating in tetraploid cell formation. Images were acquired at 5 min time resolution and timestamps are shown in hours:minutes. Scale bar, 10 µm. Related to Figure F7A-C.

**Movie S7: Ubiquitin, E4orf4, and AurB co-tracking at the midbody prior to cleavage furrow regression.**
Time-lapse imaging of AdV-C5-wt-infected cells co-expressing fluorescently tagged ubiquitin, E4orf4, and AurB. During late cytokinesis, all three signals accumulate at the midbody and display coordinated spatial and temporal dynamics. Persistent midbody association of ubiquitin, E4orf4, and AurB and removal precedes cleavage furrow regression and midbody collapse, resulting in tetraploid cell formation. Images were acquired at 15min time intervals and timestamps are shown as hours:minutes. Scale bar, 10 µm.

#### Supplementary Tables

Table S1: Proteins identified in mass spectrometry with corresponding fold changes and p-values (related to Fig. 3C).

**Key Resources Table or Table of reagents**

| REAGENT or RESOURCE | SOURCE | IDENTIFIER |
|---|---|---|
| **Antibodies and Fluorescent reagents** | | |
| Anti-proteinVI | Burckhardt et al. 2011[1] | https://doi.org/10.1016/j.chom.2011.07.006 |
| Anti-Flag | Sigma Aldrich | F7425 |
| Anti-CDC20 | Santa Cruz Biotech | sc-13162 |
| Anti-GFP | ThermoFisher | GF28R |
| Anti-AuroraB | BD Biosciences | 611083 |
| Anti-LaminA | Santa Cruz Biotech | sc-7292 |
| Anti-β-Actin | Sigma Aldrich | A5441 |
| Anti-HA | ThermoFisher | PA1-985 |
| Anti-GAPDH | ThermoFisher | PA1-987 |
| Anti-E1A (M58) | Santa Cruz Biotech | Sc-58658 |
| Alexa Fluor 488 donkey anti-rabbit IgG | ThermoFisher | A-21206 |
| Alexa Fluor 488 donkey anti-mouseIgG | ThermoFisher | A-21202 |
| Alexa Fluor 568 donkey anti-rabbit IgG | ThermoFisher | A-10042 |
| Alexa Fluor 568 donkey anti-mouse IgG | ThermoFisher | A-10037 |
| DAPI | MoBiTec | MFPCCFA-211 |
| Hoechst 33342 | Sigma Aldrich | B2261 |
| Alexa Fluor Plus 647 Wheat Germ Agglutinin | ThermoFisher | W32466 |
| Lipidtox | ThermoFisher | H34477 |
| **Pharmacological inhibitors and compounds** | | |
| Okadaic acid | Sigma Aldrich | O9381 |
| Apcin | Sigma Aldrich | SML1503 |
| ProTAME | Sigma Aldrich | SML3977 |
| Thymidine | Sigma Aldrich | T9250 |
| **Experimental models: Cell lines** | | |
| HEK293T | ATCC | #CRL-3216 |

| REAGENT or RESOURCE | SOURCE | IDENTIFIER |
|---|---|---|
| HeLa-AuroraB-EGFP-mRFP-LAP2β | Steigemenn et al. 2009[2] | https://doi.org/10.1016/j.cell.2008.12.020 |
| HeLa ATCC | This manuscript | N/A |
| HeLa-H2B-mCherry | | |
| HeLa FUCCI | Sakaue-Sawano et al. 2008[3] | https://doi.org/10.1016/j.cell.2007.12.033 |
| HeLa FUCCI-CA | Sakaue-Sawano et al. 2017[4] | https://doi.org/10.1016/j.cell.2007.12.033 |
| HeLa-Ub-mCherry | This manuscript | N/A |
| HeLa-Ub-BFP-AurB-mScar | This manuscript | N/A |
| A549-ATCC | American Type Cell Culture | Cat #CCL-185 |
| A549-sec61ß-GFP | Prasad et al. 2023[5] | https://doi.org/10.1016/j.molcel.2023.06.020 |
| A549-AurB-mScarlet | This manuscript | N/A |
| A549-CDC20-mScarlet | This manuscript | N/A |
| HEK293T-AurB-mScarlet (knock-in) | This manuscript | N/A |
| A549-Ub-GFP-AurB-mScarlet | This manuscript | N/A |
| HDF-TERT | Jing Yu et al. 2001[6] | https://doi.org/10.1006/viro.2001.1204 |
| Primary human bronchial airway basal cells | Hubert et al. 2006[7] | Donor ID: AB0839 |
| **Virus strains** | | |
| AdV-C5wt | Prasad et al. 2014[8] | http://dx.doi.org/10.1128/JVI.02156-14 |
| AdV-C2wt | Prasad et al. 2014[8] | http://dx.doi.org/10.1128/JVI.02156-14 |
| AdV-C5-GFP-E4orf4 | Miron et al. 2009[9] | https://doi.org/10.1128/JVI.01703-08 |
| AdV-C5-Flag-E4orf4 | Miron et al. 2009[9] | https://doi.org/10.1128/JVI.01703-08 |
| AdV-C5-ΔE4orf4 | Miron et al. 2009[9] | https://doi.org/10.1128/JVI.01703-08 |
| AdV-dE1-HA-E4orf4 | Miron et al. 2009[9] | https://doi.org/10.1128/JVI.01703-08 |

| REAGENT or RESOURCE | SOURCE | IDENTIFIER |
|---|---|---|
| AdV-C5-E4orf4-R81F84A | Miron et al. 2009[9] | https://doi.org/10.1128/JVI.01703-08 |
| **Oligonucleotides** | | |
| siRNA pooled oligos targeting CDC20 (NM_001255) | Dharmacon | #J-003225 |
| **Plasmids** | | |
| pCRIS-PITChv2-AurB | This manuscript | N/A |
| pLVX-CDC20-HA-IRES-Puro | This manuscript | N/A |
| pLVX-CDC20-mScarlet-IRES-Puro | This manuscript | N/A |
| pET28a-10xHis-Flag-E4orf4 | This manuscript | N/A |
| pET28a-10xHis-E1A-WT | This manuscript | N/A |
| pET28a-10xHis-CDC20(110-499)-HA-GST | This manuscript | N/A |
| pLVX-AurB-mScarlet-IRES-Puro | This manuscript | N/A |
| pLVX-AurB-HA-IRES-Puro | This manuscript | N/A |
| pLVX-GFP-Ub-IRES-Puro | Bauer et al. 2019[10] | https://doi.org/10.1016/j.celrep.2019.11.064 |
| AdV-C5-pE4-E4-GFP-E4orf4-WT | This manuscript | N/A |
| AdV-C5-pE4-E4-GFP-ΔE4orf4 | This manuscript | N/A |
| pLVX-His-Ub-IRES-Puro | This manuscript | N/A |
| **Deposited Data** | | |
| Proteomics data | PRIDE proteomics database | PXD073585 |
| **Software and Algorithms** | | |
| Analysis code for proteomics data analysis | | |
| ilastik | Berg et al., 2019[11] | https://www.ilastik.org |
| FIJI | | https://imagej.nih.gov/ij/ |
| CellProfiler | Broad Institute, USA | https://cellprofiler.org |
| CellProfiler Analyst | Broad Institute, USA | https://cellprofileranalyst.org |
| IMOD ver 4.10.42 | Kremer et al., 1996[12] | https://bio3d.colorado.edu/imod/ |

| REAGENT or RESOURCE | SOURCE | IDENTIFIER |
|---|---|---|
| Serial EM | Mastronarde, 2005[13] | https://bio3d.colorado.edu/SerialEM/ |

#### Resource Availability

##### Lead contact

##### Materials availability

Plasmids and/or cell lines will be distributed under the terms of a material transfer agreement.

#### References


1. Burckhardt, C.J., Suomalainen, M., Schoenenberger, P., Boucke, K., Hemmi, S., and Greber, U.F. (2011). Drifting motions of the adenovirus receptor CAR and immobile integrins initiate virus uncoating and membrane lytic protein exposure. Cell Host Microbe *10*, 105–117. https://doi.org/10.1016/j.chom.2011.07.006.
2. Steigemann, P., Wurzenberger, C., Schmitz, M.H.A., Held, M., Guizetti, J., Maar, S., and Gerlich, D.W. (2009). Aurora B-Mediated Abscission Checkpoint Protects against Tetraploidization. Cell *136*, 473–484. https://doi.org/10.1016/j.cell.2008.12.020.
3. Sakaue-Sawano, A., Kurokawa, H., Morimura, T., Hanyu, A., Hama, H., Osawa, H., Kashiwagi, S., Fukami, K., Miyata, T., Miyoshi, H., et al. (2008). Visualizing spatiotemporal dynamics of multicellular cell-cycle progression. Cell *132*, 487–498. https://doi.org/10.1016/j.cell.2007.12.033.
4. Sakaue-Sawano, A., Yo, M., Komatsu, N., Hiratsuka, T., Kogure, T., Hoshida, T., Goshima, N., Matsuda, M., Miyoshi, H., and Miyawaki, A. (2017). Genetically Encoded Tools for Optical Dissection of the Mammalian Cell Cycle. Mol. Cell *68*, 626-640.e5. https://doi.org/10.1016/j.molcel.2017.10.001.
5. Prasad, V., Cerikan, B., Stahl, Y., Kopp, K., Magg, V., Acosta-Rivero, N., Kim, H., Klein, K., Funaya, C., Haselmann, U., et al. (2023). Enhanced SARS-CoV-2 entry via UPR-dependent AMPK-related kinase NUAK2. Mol. Cell *83*, 2559–2577. https://doi.org/10.1016/j.molcel.2023.06.020.
6. Yu, J., Boyapati, A., and Rundell, K. (2001). Critical role for SV40 small-t antigen in human cell transformation. Virology *290*, 192–198. https://doi.org/10.1006/viro.2001.1204.
7. Hubert, M., Haemmerli, P., Marcourt, L., Lara-Quintero, E., Arthaud, L., Quiros-Guerrero, L.-M., Donnaray, S., Rimensberger, K., Gaudry, A., Alessandri-Gradt, E., et al. (2026). Human airway organoids as a high-throughput screening platform for antiviral natural products discovery. Preprint at bioRxiv, https://doi.org/10.64898/2026.02.04.703746 https://doi.org/10.64898/2026.02.04.703746.
8. Prasad, V., Suomalainen, M., Pennauer, M., Yakimovich, A., Andriasyan, V., Hemmi, S., and Greber, U.F. (2014). Chemical Induction of Unfolded Protein Response Enhances Cancer Cell Killing through Lytic Virus Infection. J. Virol. *88*, 13086–13098. https://doi.org/10.1128/JVI.02156-14.
9. Miron, M.-J., Blanchette, P., Groitl, P., Dallaire, F., Teodoro, J.G., Li, S., Dobner, T., and Branton, P.E. (2009). Localization and importance of the adenovirus E4orf4 protein during lytic infection. J. Virol. *83*, 1689–1699. https://doi.org/10.1128/JVI.01703-08.
10. Bauer, M., Flatt, J.W., Seiler, D., Cardel, B., Emmenlauer, M., Boucke, K., Suomalainen, M., Hemmi, S., and Greber, U.F. (2019). The E3 Ubiquitin Ligase Mind Bomb 1 Controls Adenovirus Genome Release at the Nuclear Pore Complex. Cell Rep *29*, 3785-3795 e8. https://doi.org/10.1016/j.celrep.2019.11.064.
11. Berg, S., Kutra, D., Kroeger, T., Straehle, C.N., Kausler, B.X., Haubold, C., Schiegg, M., Ales, J., Beier, T., Rudy, M., et al. (2019). ilastik: interactive machine learning for (bio)image analysis. Nat Methods *16*, 1226–1232. https://doi.org/10.1038/s41592-019-0582-9.
12. Kremer, J.R., Mastronarde, D.N., and McIntosh, J.R. (1996). Computer visualization of three-dimensional image data using IMOD. J Struct Biol *116*, 71–76. https://doi.org/10.1006/jsbi.1996.0013.
13. Mastronarde, D.N. (2005). Automated electron microscope tomography using robust prediction of specimen movements. J Struct Biol *152*, 36–51. https://doi.org/10.1016/j.jsb.2005.07.007.
